# STELLA: Towards a Biomedical World Model with Self-Evolving Multimodal Agents

**DOI:** 10.1101/2025.07.01.662467

**Authors:** Ruofan Jin, Mingyang Xu, Fei Meng, Guancheng Wan, Qingran Cai, Yize Jiang, Jin Han, Yuanyuan Chen, Wanqing Lu, Mengyang Wang, Zhiqian Lan, Yuxuan Jiang, Junhong Liu, Dongyao Wang, Le Cong, Zaixi Zhang

## Abstract

The staggering complexity of modern biomedical research has intensified the aspiration for a generalist “Biomedical World Model”, yet current AI agents remain constrained by static capabilities and a lack of self-evolution mechanisms. To bridge this gap, we present STELLA, a self-evolving multimodal agent designed to progressively refine its computational reasoning and physical execution through interaction. STELLA operates via a collaborative multi-agent framework (comprising Manager, Developer, Critic, Critic, and Tool Creation agents) that continuously updates reasoning templates and autonomously expands a dynamic “Tool Ocean”. We demonstrate STELLA’s capabilities on the created Tool Creation Benchmark, where it attains a score of 4.01/5 with 100% task completion, significantly outperforming state-of-the-art models including GPT-5, Claude 4 Opus, and Biomni. Beyond computational metrics, STELLA drives experimentally validated scientific discovery. In oncology, the agent identified Butyrophilin Subfamily 3 Member A1 (BTN3A1) as a novel negative regulator of natural killer (NK) cell function in acute myeloid leukemia (AML), verified via CRISPR knockout studies. In protein engineering, STELLA orchestrated a complete directed evolution workflow for the enzyme strictosidine synthase, identifying variants, notably M276L, exhibiting more than a two-fold improvement in catalytic activity. Finally, the system extends to physical laboratory automation by training Vision-Language-Action (VLA) models through a Decompose-Monitor-Recover mechanism, which increased success rates from 17% to 82%. By integrating autonomous tool evolution, biological discovery, and robotic control, STELLA offers a blueprint for a self-evolving world model in the life sciences.

## Introduction

Modern biomedical research is defined by a dual reality: immense opportunity coupled with staggering complexity, driven by rapidly expanding datasets, heterogeneous experimental modalities, and the rapid acceleration of methodology development^1, 2^. This landscape has intensified the long-standing aspiration for a Biomedical World Model—a unified, self-evolving representation that integrates multimodal knowledge, predictive modeling, and physical interaction to support autonomous scientific discovery^3–6^. In this work, we present STELLA to advance this goal. STELLA is a self-evolving multimodal agent designed to seamlessly integrate computational reasoning with wet-lab verification. As illustrated in Fig. 1a, STELLA engages with research tasks by utilizing reasoning templates for memory, bio-foundation models as tools, and actions ranging from coding to robotic control. Through this process, feedback from genes, cells, and wet-lab experiments continually updates its internal representations, facilitating the formation of a world model grounded in real-world verification.

**Figure 1.**
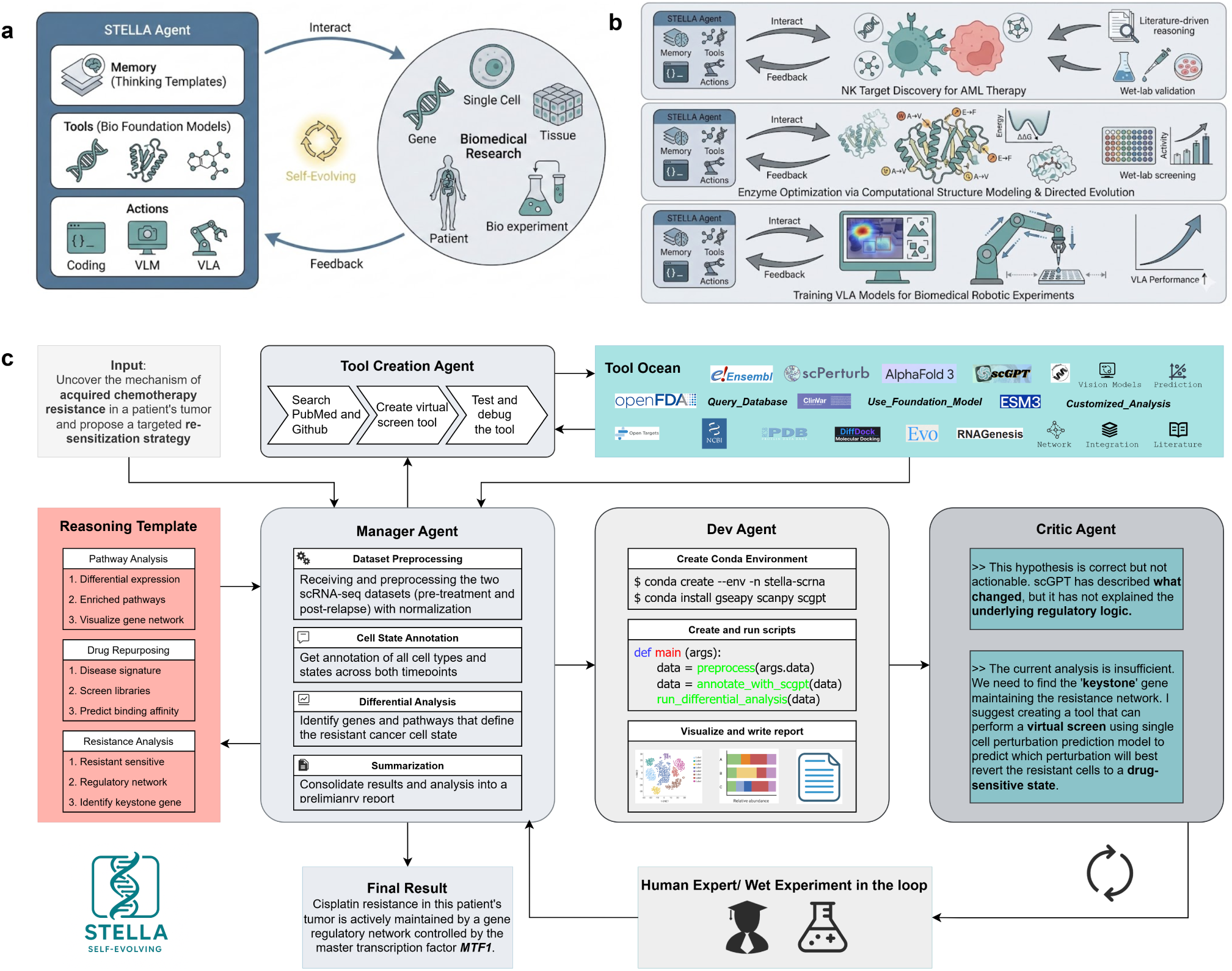
STELLA: A self-evolving multimodal agent system towards biomedical world model. **a,** Conceptual view of how STELLA interacts with the biomedical research environment. STELLA uses memory (reasoning templates), a suite of biological models as tools, and multimodal actions, including coding, Vision-Language Models (VLMs), and Vision-Language-Action (VLA) models to engage with genes, cells, tissues, patients, and wet lab experiments. Feedback from these interactions updates STELLA’s internal templates and tools and facilitates the formation of a biomedical world model. **b,** Representative applications. Top: discovery of natural killer (NK) cell targets for acute myeloid leukemia (AML) therapy through literature reasoning and wet-lab validation. Middle: enzyme optimization supported by computational structure modeling and directed evolution experiments. Bottom: training of VLA models for robotic execution of biomedical procedures and improvement of manipulation performance. **c,** Internal architecture of STELLA. Reasoning templates guide the Manager Agent through data preprocessing, annotation, analysis and summarization. The Dev Agent executes computational workflows. The Critic Agent evaluates hypotheses and proposes new analyses or tool creation needs. The Tool Creation Agent continuously expands the Tool Ocean with new computational tools. Human and experimental feedback closes the loop, enabling self-evolution across tasks.

The distinguishing feature of STELLA is its ability to transcend static automation through two novel self-evolution mechanisms that allow it to become more proficient with experience^7, 8^. First, rather than reinitiating the process for every task, STELLA maintains a Template Library of reasoning workflows. This library is dynamically updated with successful strategies, allowing the system to distill generalizable logic from specific cases and progressively refine its problem-solving patterns. Second, we replace the conventional fixed toolkit with a Tool Ocean—a dynamic and growing repository of bioinformatics resources. Unlike static systems constrained by pre-installed packages, STELLA is equipped to autonomously identify, validate, and integrate new tools and APIs in response to evolving research demands. Together, these mechanisms ensure that STELLA’s knowledge base and technical capabilities continuously expand as it interacts with the scientific environment.

To implement these evolutionary mechanisms, STELLA employs a collaborative multi-agent framework comprising four specialized agents (Manager, Developer, Critic, and Tool Creation). The workflow begins with the Manager Agent, which retrieves relevant strategies from the Template Library to coordinate a multi-step reasoning plan. The Developer Agent then executes these steps by generating and running Python scripts for bioinformatics analyses. Crucially, this process is governed by a rigorous feedback loop: the Critic Agent continuously assesses intermediate results to identify flaws, prompting the Developer to refine its code until the objective is achieved^9, 10^. Meanwhile, the Tool Creation Agent operates in the background to populate the Tool Ocean, actively discovering and wrapping new tools required by the current workflow. This architecture ensures that STELLA does not merely execute commands, but engages in a robust, feedback-driven cycle of reasoning, execution, and self-correction.

To assess STELLA’s capabilities, we first evaluated it on rigorous public benchmarks. On the challenging Humanity’s Last Exam Biomedicine benchmark^11^, STELLA achieved an accuracy of 32%, surpassing all state-of-the-art Large Language Models (including GPT-5^12^, Gemini 2.5 Pro^13^, and DeepSeek R1^14^) and the specialized agents (including Biomni^15^ and ToolUniverse^16^). Similarly, on the LAB Bench suite^17^, STELLA demonstrated strong generalization with top-tier performance on both DBQA (61%) and LitQA (65%). To further probe the capacity for autonomous tool use, we introduced the **Tool Creation Benchmark**, which comprises 47 tasks across 10 biomedical domains. STELLA reached the highest mean score of 4.01/5 while maintaining a 100% task completion rate. This performance establishes a substantial lead over all baselines: the nearest competitor, the specialized agent Biomni, achieved a score of 3.33/5 despite also maintaining 100% completion. Generalist Large Language Models (LLMs) achieved scores lower than 3.0 (e.g., Gemini 2.5 Pro achieved 2.99/5 with an 85.1% completion rate). These results highlight the advantage of a self-evolving agent that integrates tool creation, computational analysis, and language-driven reasoning for complex biomedical tasks.

Beyond standardized evaluation, a genuine biomedical world model must demonstrate the capacity to generate novel insights amenable to wet-lab verification. To validate this predictive capability, we first applied STELLA to **identify previously unreported negative regulators of natural killer (NK) cell function in acute myeloid leukemia (AML)**. By synthesizing data from molecular databases, literature, and regulatory networks, STELLA prioritized **Butyrophilin Subfamily 3 Member A1 (BTN3A1)** as the most promising candidate for therapeutic targeting. CRISPR (Clustered Regularly Interspaced Short Palindromic Repeats) knockout studies in the NK92MI cell line experimentally validated this finding. Loss of BTN3A1 consistently increased NK cell cytotoxicity across four leukemia models and enhanced degranulation as measured by CD107a expression. These results demonstrate that STELLA can autonomously propose hypotheses that lead to validated biological findings, highlighting its ability to uncover therapeutically relevant mechanisms.

We further examined whether STELLA could achieve closed-loop optimization between computational design and wet-lab verification by supporting **multi-round enzyme engineering**. Using strictosidine synthase as a target system, STELLA generated a complete directed evolution workflow that integrated structural modeling, virtual mutagenesis, activity prediction (via a fine-tuned protein language model), and stability filtering. Two iterative rounds of design and experimental testing identified three mutations with markedly increased catalytic activity. Notably, the **M276L variant improved product formation by more than two-fold**, while the V176F and M276R variants also displayed significant improvements. Structural analysis attributed these enhancements to strengthened hydrogen bonding and a more preorganized catalytic pocket, providing a mechanistic basis for the observed improvements. These wet-lab results validate the generality of STELLA’s design agent and confirm its ability to drive experimentally actionable enzyme optimization.

Finally, to establish the biomedical world model’s physical grounding, STELLA extends beyond computational tasks to coordinate complex laboratory workflows through Vision-Language-Action (VLA) models. Biomedical robotics presents unique challenges involving heterogeneous interfaces, ambiguous materials, deformable containers, and strictly constrained procedures that demand high precision^18–20^. To address these, STELLA adopts a **Decompose-Monitor-Recover** mechanism that segments long-horizon tasks into interpretable subtasks, continuously monitors execution using multimodal perception, and invokes specialized tools to recover from failures. We evaluated this approach on the Autobio benchmark^21^, encompassing tasks requiring liquid handling, dual-arm coordination, and contact-rich tube placement. STELLA consistently improved success rates with accumulated recovery experience, significantly outperforming supervised fine-tuning (SFT) and online reinforcement learning (RL) baselines across multiple base architectures (*π*_0_^19^, *π*_0.5_^20^, and RDT^18^). For example, success on the *π*_0_ backbone increased from 17% to 82%. These results demonstrate the critical role of active, self-evolving feedback loops in developing physical intelligence, marking a crucial step toward a biomedical world model grounded in real-world interaction.

## Results

### STELLA’s Self-evolving Mechanisms for Biomedical Research

A defining feature of STELLA is its dual **self-evolving mechanism**, which empowers it to learn from experience and continuously expand its abilities as a **Biomedical World Model** (Fig. 1a). The first mechanism involves the evolution of its **Template Library**. STELLA captures successful multi-step workflows, such as the progression from initial descriptive analysis to a predictive virtual screen, and abstracts them into high-quality reasoning templates. This process refines STELLA’s strategic knowledge, enabling it to solve similar problems more efficiently in the future.

The second, more profound level of evolution is the expansion of the **Tool Ocean**, a dynamic repository of STELLA’s executable capabilities. This ocean contains a diverse array of computational tools broadly classified into three main categories: (1) functions for querying established scientific databases, (2) interfaces for leveraging large-scale foundation models, and (3) customized analysis tools. The first category provides direct access to critical repositories like `PubMed`^22^, `ClinVar`^23^, and `PDB`^24^. The second allows STELLA to harness state-of-the-art AI, including `AlphaFold3`^25^ for structure prediction, `scGPT`^26^ for single-cell interpretation, and `ESM3`^27^ for protein language modeling. The third category consists of specialized scripts (e.g., for network analysis) and interfaces for wet-lab protocol orchestration and VLA-based real-world interaction. Together, the refinement of the Template Library and the expansion of the Tool Ocean serve as the foundational engines enabling STELLA to navigate scientific complexity with growing sophistication.

In the following sections, we will detail how STELLA’s self-evolving framework powers diverse biomedical discovery (Fig. 1b). First, we present the discovery of novel NK cell targets for AML through computational reasoning and wet-lab validation. Second, we highlight enzyme optimization achieved through iterative computational modeling and experimental feedback. Finally, we show the tool-augmented training mechanism of STELLA-VLA for self-improving robotic control, marking one initial step towards physical intelligence.

### STELLA’s Overall Framework

To operationalize these self-evolving mechanisms, STELLA leverages four key agents (Manager Agent, Dev Agent, Critic Agent, and Tool Creation Agent) to systematically address complex biomedical research questions (Fig. 1c). The workflow begins when the **Manager Agent** receives a high-level research goal, such as to “*uncover the mechanism of acquired chemotherapy resistance and propose a re-sensitization strategy.*” The Manager Agent analyzes this goal and, guided by its reasoning experience (retrieved from the Template Library), establishes a “Reasoning Pathway”—a strategic plan decomposing the problem into steps like ‘Differential expression analysis’ and ‘Identify keystone gene’. It assigns initial tasks to the **Dev Agent**, the computational workhorse, which creates a self-contained conda environment and executes analysis scripts (e.g., `diff analysis.py`). The results are passed to the **Critic Agent** for rigorous evaluation. In the chemoresistance example, the Critic might provide feedback such as: “*This hypothesis is correct but not actionable… It describes **what** changed but not the regulatory logic. We need to find the ‘keystone’ gene.*” This feedback identifies a capability gap. In response, the Manager Agent tasks the **Tool Creation Agent** to close this gap. This agent leverages the **Tool Ocean** described above to build or configure a new tool, such as a virtual perturbation screening model based on virtual cell states^28^. By deploying this new tool, STELLA moves beyond simple description to mechanistic interpretation, ultimately identifying the transcription factor `MTF1` as the regulator of the resistance network. Currently, STELLA utilizes Claude 4 Sonnet for the Dev and Tool Creation Agents, and Gemini 2.5 Pro for the Manager and Critic Agents.

### Benchmarking Tool Creation and Tool-Augmented Biomedical Reasoning

To systematically benchmark STELLA and leading LLMs/agent systems in domain-appropriate tool creation and tool-augmented biomedical research, we developed the **Tool Creation Benchmark**. The benchmark is generated through a structured six-stage workflow that captures how real scientific tasks arise in biomedical research (Fig. 2a). The process begins with identifying core biomedical domains and their associated experimental, computational, and database-driven tools, before collecting high-quality task scenarios from recent research articles. These scenarios reflect authentic scientific operations such as pathway-to-gene mapping, protein chain-level analysis, and primer design under thermodynamic constraints. Domain experts then refine each scenario into a precise task prompt with explicit tool intent, clearly defined inputs and outputs, and guidelines that encourage tool-grounded reasoning. Expected answers are specified in structured formats, with validated reference outputs or accepted tool results. The final benchmark contains 47 tasks spanning 10 biomedical fields, from molecular biology and protein design to pharmacology and synthetic biology, as illustrated in Fig. 2b & c. Evaluation uses a five-dimensional rubric that scores technical accuracy, domain knowledge, analytical reasoning, tool innovation, and communication clarity on a 1–5 scale. This evaluation design not only measures whether a system produces the correct answer, but also whether it does so through transparent, reproducible, and tool-grounded reasoning. The tool creation benchmark aims to expose the gap between static LLM knowledge and the dynamic, tool-grounded reasoning required for genuine scientific discovery.

**Figure 2.**
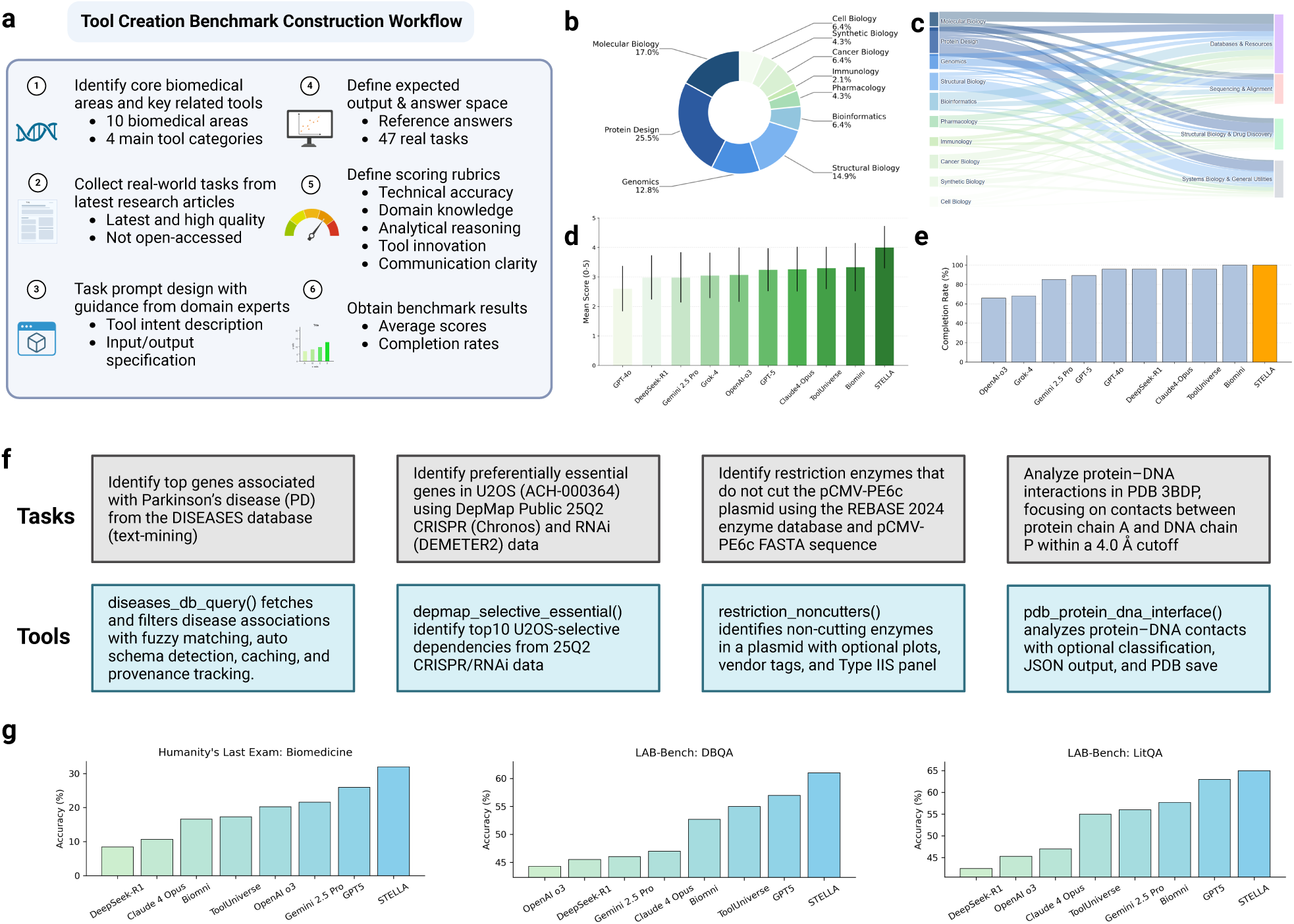
Benchmarking tool creation capability of STELLA. **a**, Workflow for constructing the tool creation benchmark, which integrates ten biomedical domains and four major tool categories, with 47 real-world tasks from recent research articles. **b–c**, Distribution of domains and task types across the benchmark. **d–e**, Overall performance of baseline models and STELLA on tool creation, reported as mean scores (**d**) and task completion rates (**e**). **f**, Examples of novel tools autonomously created by STELLA, including disease gene association queries, selective essentiality analysis from CRISPR/RNAi data, non-cutting enzyme identification, and protein–DNA interface analysis. **g**, Comparative evaluation of STELLA against frontier LLMs and agents on widely used external benchmarks (Humanity’s Last Exam: Biomedicine, DBQA, and LitQA), demonstrating superior accuracy in diverse biomedical reasoning tasks.

In Fig. 2d & e, we compared STELLA with biomedical agents (Biomni^15^ and ToolUniverse^16^) and seven leading LLMs (GPT o3^29^, GPT-4o^30^, Claude 4 Opus^31^, GPT-5^12^, Grok-4^32^, Gemini 2.5 Pro^13^, and DeepSeek R1^14^) using the same tasks and scoring rubric. STELLA achieved the highest mean score of 4.01 out of 5 with 100% task completion, outperforming all baselines. Biomni and ToolUniverse achieved a mean score of 3.33 and 3.30, respectively. Among LLMs, Claude 4 Opus reached 3.26 (100% completion), GPT-5 reached 3.24 (93.6%), GPT o3 reached 3.08 (70.2%), Grok-4 reached 3.05 (70.2%), Gemini 2.5 Pro reached 2.99 (89.4%), DeepSeek R1 reached 2.98 (100%), and GPT-4o reached 2.60 (100%). While baseline models demonstrate strong fluency and factual recall, they frequently struggle with tasks demanding complex tool execution, structured data parsing, and multi-step reasoning. These limitations are particularly evident in workflows involving highly customized database queries. For instance, ToolUniverse performs well when leveraging its pre-existing tool library but lacks the flexibility to generalize to unseen tasks. In contrast, STELLA excels in high-complexity scenarios requiring tool composition, API invocation, chained logic, and robust error recovery, consistently delivering correct, transparent, and tool-grounded outputs.

In experiments, we observed that STELLA solves benchmark tasks by dynamically selecting and executing a wide range of domain-specific tools, including molecular databases such as `KEGG`^33^, `DISEASES`^34^, and `STRING`^35^; sequencing utilities such as `BWA MEM`^36^ and `MACS2`^37^; molecular-cloning tools such as `Primer3`^38^ and `NEBcloner`^39^; and structural-biology software such as `PyMOL`^40^ and `AlphaFold2`^41^. Fig. 2f highlights several examples of novel tools autonomously created by STELLA, including disease–gene association modules, selective-essentiality analysis from CRISPR and RNA interference (RNAi) datasets, non-cutting enzyme identification, and protein–DNA interface analysis, with further details provided in Fig. S1. STELLA’s strong tool-creation ability establishes the foundation needed to solve complex biomedical problems, enables autonomous scientific discovery, and supports progress toward an integrated biomedical reasoning framework.

### STELLA Outperforms State-of-the-art LLMs and Agents on Popular Benchmarks

To further evaluate STELLA’s overall performance, we benchmarked it against a suite of SOTA large language models (DeepSeek-R1, Claude 4 Opus, OpenAI o3, Gemini 2.5 Pro, and GPT5) and specialized agents (Biomni and ToolUniverse) on three challenging biomedical question-answering tasks. The results, presented in Figure 2g, show that STELLA consistently achieves superior performance across all benchmarks. On the `Humanity’s Last Exam (Biomedicine)`^11^ benchmark, STELLA achieved a top accuracy of 32%, surpassing all other tested models. This lead was extended on the `LAB-Bench`^17^ suite, where STELLA achieved the highest scores of approximately 61% on the DBQA task and 65% on the LitQA task. These results validate the efficacy of its integrated multi-agent architecture in comparison to both generalist models like Gemini 2.5 Pro^13^ and other specialized agents.

### STELLA Discovers Novel Target of NK Cell Antileukemic Activity

NK cells play a critical role in immune surveillance against AML, but their function is frequently suppressed by tumor-intrinsic and microenvironmental inhibitory signals^42, 43^. AML blasts, for instance, may overexpress inhibitory ligands such as Human Leukocyte Antigen-E (HLA-E), which engages NK-cell inhibitory receptors like Natural Killer Group 2, member A (NKG2A), dampening NK cytotoxic activity and enabling immune evasion^44^. Although immune checkpoint blockade has transformed cancer therapy—most notably with PD-1/PD-L1 inhibitors in T-cell–based immunotherapies, the development of NK-cell–directed checkpoint inhibitors remains significantly underdeveloped^45^. This gap is particularly pressing in AML, where the absence of selective tumor-specific markers complicates both CAR-T and NK-based strategies^46^. Moreover, AML relapse is frequently driven by leukemic stem cells (LSCs), which are poorly cleared by standard therapies and remain elusive to immune targeting due to antigen similarity with normal hematopoietic stem cells^47^. Consequently, identifying novel and unreported NK-cell inhibitory regulators could offer a powerful means to boost NK-mediated immunity without compromising healthy tissues. However, due to the complexity of NK receptor-ligand interactions and extensive prior research on canonical checkpoints (e.g., NKG2A, TIGIT, PD-1), uncovering previously uncharacterized regulators is a nontrivial challenge.

To systematically uncover such hidden targets, STELLA was provided with the prompt “*Identify novel, previously unreported negative regulators of NK cell function in the context of AML*” (Fig. 3a). STELLA first employs reasoning templates that integrates literature mining with database analysis to propose candidate immunoregulatory targets. In brief, it searches biomedical databases and prior studies to identify genes associated with NK-cell regulation in AML, then filters out those already reported or well-characterized, thereby focusing on candidates that have not been previously implicated (i.e., truly novel targets). Each candidate is further validated using supporting evidence from patient datasets (e.g., gene expression patterns in the TCGA AML cohort) to ensure disease relevance and provide grounded biological rationale (Fig. 3b). This automated approach greatly accelerates hypothesis generation (Fig. 3c). STELLA completed its NK target discovery pipeline in about 8 minutes, a task that required on the order of two days of intensive manual curation by a human expert (e.g., a PhD-level researcher majoring in NK therapies for AML).

**Figure 3.**
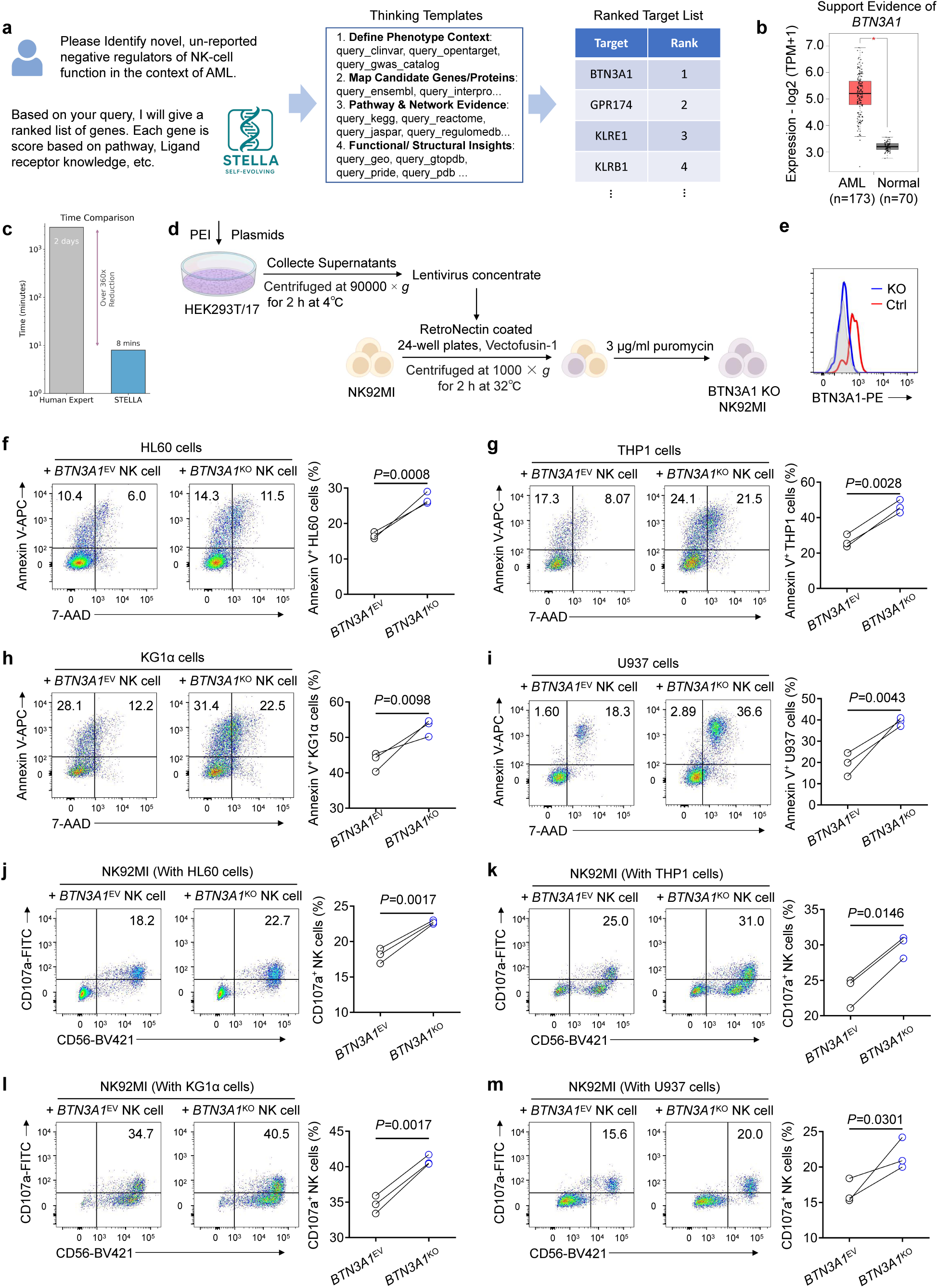
STELLA-driven discovery and validation of Butyrophilin Subfamily 3 Member A1 (BTN3A1) as a negative regulator of NK-cell anti-leukemic activity. **a–c,** STELLA identifies **BTN3A1 as a previously unreported inhibitory regulator of NK-cell function relevant to AML.** STELLA employs structured reasoning templates to parse multi-omic, pathway, and ligand–receptor knowledge, generating a ranked list of candidate negative regulators (**a**). The full target list is shown in the table. S4. BTN3A1 is supported by transcriptomic evidence from the TCGA AML dataset (**b**), where BTN3A1 expression is significantly elevated in AML samples relative to normal controls. Compared with human experts, STELLA achieves substantially accelerated target identification (**c**). **d–e**, Generation and validation of **BTN3A1^KO^ NK92MI** cells. Lentiviral transduction, RetroNectin^TM^-assisted infection, and puromycin selection yield BTN3A1-deficient NK92MI cells (**d**), with efficient knockout confirmed by flow cytometry (**e**). **f–i**, Loss of BTN3A1 enhances NK-cell cytotoxicity against AML cells. BTN3A1^KO^ NK92MI cells exhibit significantly increased killing of both TP53-wildtype (HL60) and TP53-mutant (THP-1, KG-1*⍺*, U937) AML lines, as measured by Annexin V^+^ target-cell frequency following 5 h co-culture at a 5:1 effector-to-target ratio. **j–m**, BTN3A1 knockout increases NK-cell degranulation. Across all AML co-culture settings, BTN3A1^KO^ NK92MI cells display elevated CD107a^+^ NK-cell frequencies, indicating enhanced activation and cytotoxic potential. All statistical analyses were performed using two-tailed unpaired Student’s *t*-tests. *P* < 0.05 (*), *P* < 0.01 (**), *P* < 0.001 (***), *P* < 0.0001 (****). Data are presented as means ± standard deviation.

Using STELLA, we identified **Butyrophilin Subfamily 3 Member A1** (**BTN3A1**, also known as **CD277**) as a top candidate inhibitory regulator of NK cells in AML. BTN3A1 is an immunomodulatory surface protein known for its role in T-cell biology. For example, BTN3A1 can bind to N-glycosylated CD45 and thereby inhibit T-cell receptor signaling, and it is required for certain *γδ* T-cell activation processes. There is also some evidence that BTN3A family molecules can regulate NK-cell activity (affecting NKp30-mediated IFN*γ* production). BTN3A1 is supported by transcriptomic evidence from the TCGA AML dataset organized by STELLA (Fig. 3b), where BTN3A1 expression is significantly elevated in AML samples relative to normal controls. However, prior to this work, BTN3A1 had not been reported to be linked to NK-dysfunction. STELLA’s analysis singled out BTN3A1 despite its absence from existing NK immunotherapy literature, highlighting the platform’s ability to uncover non-obvious targets. These insights prompted us to perform experimental evaluation of BTN3A1’s role in NK-cell anti-leukemia responses.

We next performed a series of in vitro experiments to functionally validate BTN3A1 as a negative regulator of NK-cell activity. We used CRISPR/Cas9-genome-editing to generate a BTN3A1 knockout in the NK92MI cell line, a well-established human NK-cell model, and confirmed the loss of BTN3A1 protein expression by flow cytometry (Fig. 3d & e). To assess cytotoxic function, we co-cultured BTN3A1^KO^ NK92MI cells with four AML cell lines. These included three TP53-mutant lines (THP-1, KG-1*⍺*, U937) and one TP53-wildtype line (HL60). Across all four models, BTN3A1-deficient NK cells demonstrated significantly increased killing of AML targets, as indicated by elevated Annexin V positivity in the target cells (Fig. 3f–i). We further examined NK-cell effector function by measuring surface CD107a expression, a marker of degranulation and cytolytic granule release. BTN3A1 knockout cells consistently showed higher CD107a levels following co-culture with AML cells (Fig. 3j–m), confirming enhanced activation. Together, these results establish BTN3A1 as a previously unrecognized inhibitory regulator of NK-cell function. Its disruption significantly enhances NK-mediated cytotoxicity and degranulation against AML, supporting BTN3A1 as a promising therapeutic target and validating STELLA’s prediction. This workflow exemplifies STELLA’s ability to condense labor-intensive hypothesis generation into rapid, high-fidelity reasoning for accelerated scientific discovery.

### STELLA Optimizes Enzyme Activity for Strictosidine Synthase

To demonstrate the potential of AI-assisted enzyme engineering in advancing biological applications, we focused on optimizing the activity of strictosidine synthase from *Rauvolfia serpentina* (*Rs*STR; UniProt ID: P68175). Terpenoid indole alkaloids (TIAs) are a structurally diverse group of natural products primarily biosynthesized by plants of the Apocynaceae family^48^. As essential biogenic precursors, TIAs form the basis for numerous clinically valuable therapeutics, including anticancer agents such as vinblastine, vincristine, and irinotecan^49, 50^. A critical, rate-limiting step in TIA biosynthesis is the *Rs*STR-catalyzed Pictet–Spengler condensation of tryptamine and secologanin to form strictosidine^51–53^. Consequently, engineering *Rs*STR variants with improved catalytic activity is crucial for enhancing strictosidine production and enabling the efficient biosynthesis of medically relevant TIAs. While previous studies have identified mutations that modulate activity, these efforts were largely empirical and limited in scope. Deep learning models, particularly when augmented by a self-evolving agent, offer a transformative approach to this challenge, accelerating the discovery of highly active enzymes through iterative rounds of directed evolution.

Prompted by the user command *“Optimize the catalytic activity of strictosidine synthase (UniProt ID: P68175) for its natural reaction involving tryptamine and secologanin,”* STELLA automatically initiated a multi-round directed evolution workflow. This process integrates structural modeling, protein language model prediction, and continuous wet-lab feedback (Fig. 4a). Drawing from prior successful workflows stored in its evolving template library, the Manager Agent defines a four-step reasoning pathway: (1) retrieval of enzyme and substrates, (2) structural modeling and catalytic pocket identification, (3) mutant library construction and activity prediction, and (4) biophysical stability filtering and experimental recommendation.

**Figure 4.**
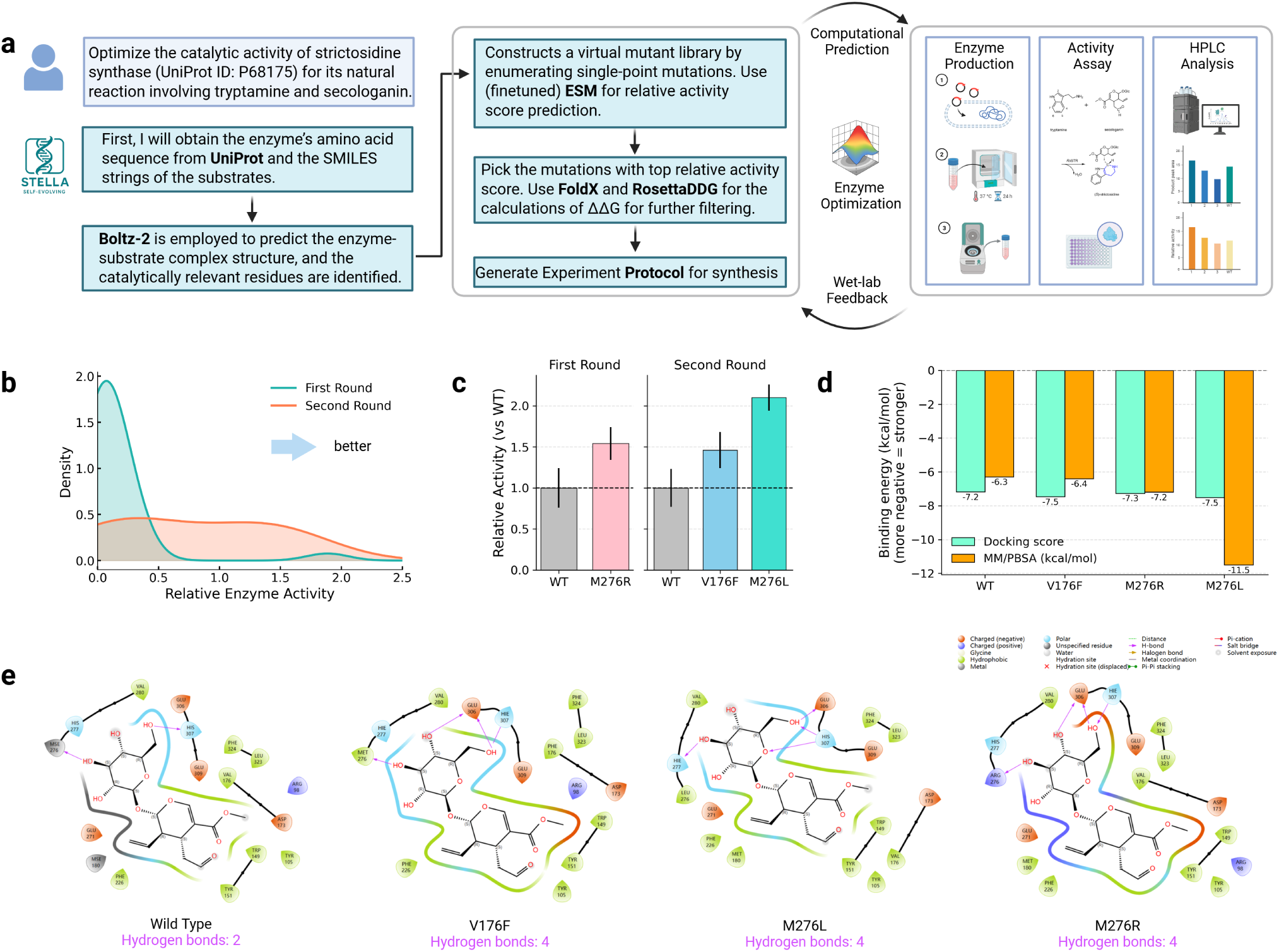
STELLA-driven multi-round enzyme optimization with wet-lab feedback. **a**, Workflow of STELLA for optimizing the catalytic activity of strictosidine synthase from *Rauvolfia serpentina* (*Rs*STR; UniProt ID: P68175), integrating computational prediction, mutational library design, and wet-lab validation. After each round, wet-lab experimental data are reintegrated to fine-tune predictive models, enabling iterative improvement. **b**, Distribution of relative enzyme activity across the first and second optimization rounds, demonstrating improved activity after feedback integration. **c**, Experimental validation of selected STR variants across two rounds. M276R was tested relative to the WT in the first round, and V176F and M276L in the second round, all of which showed enhanced catalytic activity. Each variant or WT activity was measured in three independent experiments. **d**, Computational evaluation of binding energies for WT and variants, measured by docking score and MM/PBSA calculations, correlating with increased activity in high-performing mutants. **e**, Interaction diagrams of wild type and variants highlighting hydrogen-bond networks, drawn using Schrödinger software, with variants forming more stabilizing interactions compared to WT.

Specifically, in Step 1, the Dev Agent retrieves the *Rs*STR amino acid sequence from UniProt^54^ and the SMILES representations of tryptamine and secologanin using the Tool Ocean. In Step 2, STELLA employs Boltz-2^55^ to predict the enzyme–substrate complex and identify pocket residues. Based on these pocket residues, the Dev Agent constructs a virtual single-point mutant library by enumerating all substitutions within a 5 °A radius of the active site while excluding residues essential for global folding. For each candidate mutant, STELLA predicts relative activity using a fine-tuned ESM2 (650M)^56^ protein language model (Step 3). This ESM2 model is iteratively refined through STELLA’s self-evolving mechanism: experimental data generated from the preceding round are used to retrain the model, improving its *Rs*STR-specific sequence–activity mapping and thereby yielding increasingly accurate predictions in subsequent rounds. In Step 4, STELLA further filters high-scoring mutants using thermodynamic stability criteria. FoldX^57^ and RosettaDDG^58^ are invoked to compute ΔΔ*G* values, and only variants with favorable stability (typically ΔΔ*G* < 0 kcal/mol in both models) pass the filter. The Critic Agent then evaluates mechanistic plausibility, flags potential structural risks, and suggests refinements prior to generating the final set of recommended mutants and the associated experimental protocol for synthesis.

In the first round, the workflow generated 30 single-point variants targeting the catalytic pocket. Although most mutants showed reduced activity relative to wild type (WT), several displayed moderate improvements, yielding a valuable sequence–function dataset. Notably, M276R improved activity by 54%, highlighting Met276 as a key hotspot for optimization. STELLA automatically incorporated these measurements to fine-tune the ESM2 model, substantially enhancing its predictive accuracy for *Rs*STR.

In the second round, candidate mutations were re-ranked using the updated ESM2 model, structural stability filters (FoldX and RosettaDDG), and STELLA’s Critic Agent to evaluate mechanistic plausibility. STELLA subsequently recommended a prioritized panel of variants for experimental testing. For example, **M276L increased catalytic yield by 110%** compared to WT. Density distributions across rounds confirm a clear rightward shift in activity (Fig. 4b), demonstrating that STELLA effectively incorporates wet-lab feedback to progressively enrich for high-performing mutants. Docking scores and MM/PBSA calculations of high-performing mutants are also higher than WT, demonstrating consistency with wet-lab activity (Fig. 4d). Structural interaction diagrams reveal strengthened hydrogen-bond networks and more favorable substrate–enzyme contacts in the optimized variants (Fig. 4e), offering a mechanistic basis for their improved kinetics.

To provide a mechanistic explanation for the experimentally observed catalytic gains, STELLA also autonomously orchestrated molecular dynamics (MD) simulations to characterize the structural stability and dynamic behavior of the top-performing variants. By coordinating the analysis of backbone Root Mean Square Deviation (RMSD), per-residue Root Mean Square Fluctuation (RMSF), and the radius of gyration (*R*_g_) over production trajectories (Fig. S4, S5, and S6), the agent correlated conformational dynamics with functional improvements. For each system, replicate simulations were averaged and visualized with standard deviation bands. STELLA observed that backbone RMSD traces for all variants reached stable plateaus following equilibration (Fig. S4), confirming the maintenance of global structural integrity comparable to WT. Similarly,

**Figure 5.**
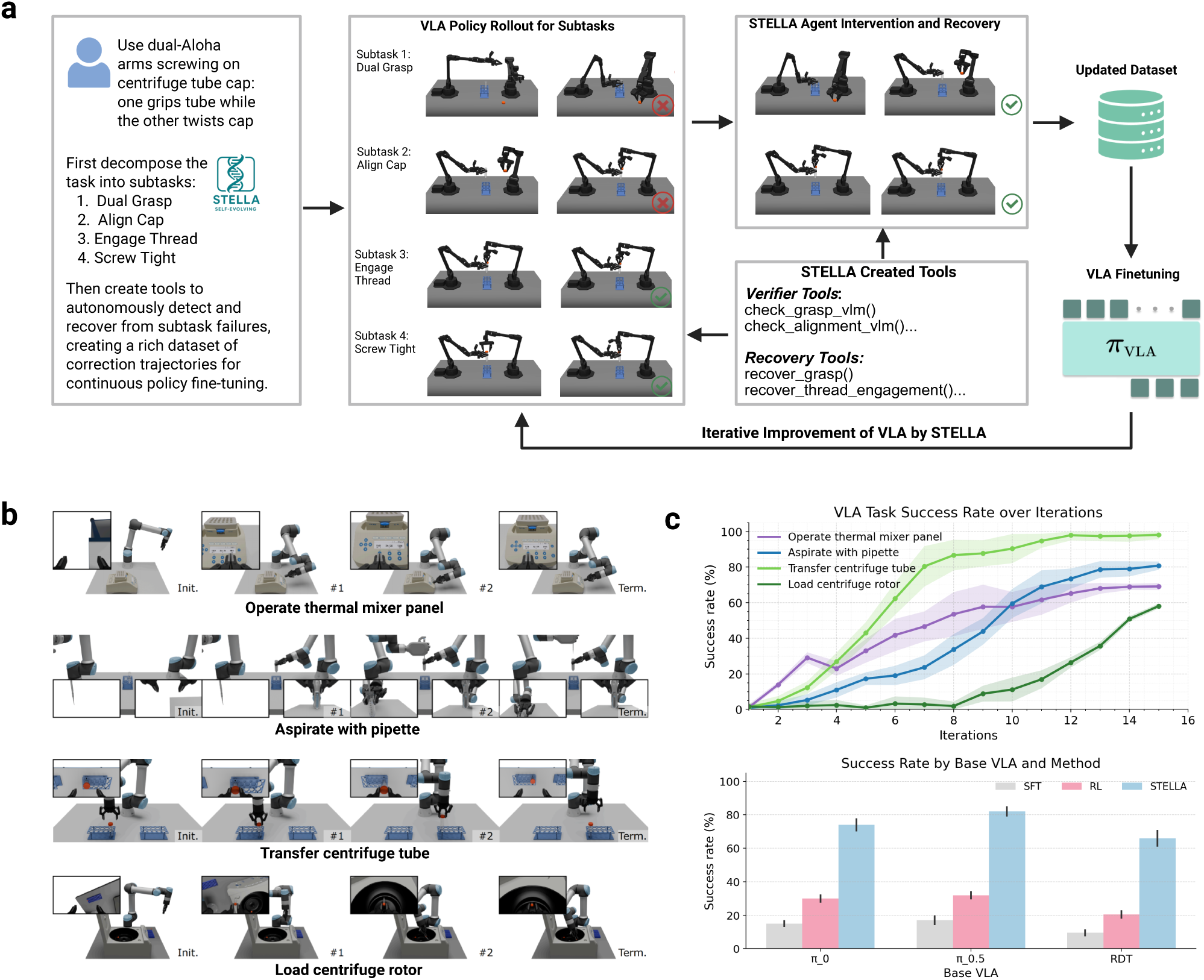
STELLA-driven physical intelligence built up through self-evolving Vision-Language-Action (VLA). **a**, The framework employs a “Decompose-Monitor-Recover” mechanism. Unlike static training, STELLA actively creates tools to interact with the environment when errors occur. This process generates a rich dataset of recovery trajectories, effectively teaching the VLA the causal dynamics of the physical world (i.e., how to recover from failure). **b**, Diverse biomedical tasks conducted by STELLA-VLA. Visualizations of four representative long-horizon tasks ranging from precise liquid-handling (Aspirate with pipette) to contact-rich manipulation (Load centrifuge rotor). **c**, Self-improvement and comparison. Top: STELLA-VLA’s success rate steadily increases over iterations, demonstrating the accumulation of world knowledge. Shaded regions represent the standard deviation across 5 independent runs. Bottom: STELLA-VLA significantly outperforms traditional SFT and RL baselines across different VLA architectures (*π*_0_, *π*_0.5_, RDT) with an equivalent sample budget, validating the efficacy of tool-mediated evolution. Error bars indicate the standard deviations.

RMSF profiles revealed only localized changes in flexibility near mutated regions, with global fluctuation patterns largely preserved (Fig. S5). Crucially, STELLA identified a consistent decrease in *R*_g_ values for the optimized variants compared to WT (Fig. S6), indicating a modestly more compact conformational ensemble. From this multimodal data, STELLA concluded that this reduced structural heterogeneity, combined with preserved stability, fosters enhanced active-site preorganization—a key dynamic mechanism driving the improved catalytic activity.

### STELLA-Driven Physical Intelligence for Biomedical World Modeling

VLA architectures jointly integrate visual observations, natural language instructions, and proprioceptive cues to generate action trajectories in an end-to-end manner^18–20^. By aligning perception, reasoning, and control more closely with how humans interact with the physical world, VLA systems represent a promising foundation for developing generalist world models that unify multimodal understanding with embodied action. In STELLA, we aim to specifically advance biomedical experimental robotics, which presents unique challenges because robots must operate across heterogeneous laboratory interfaces such as digital displays, rotating knobs, membrane buttons, tubes, and transparent containers while maintaining strict millimeter-level precision. The presence of visually ambiguous materials, deformable vessels, and tightly constrained multi-step operations further demands strong visual reasoning, reliable language understanding, and consistently high-precision manipulation that exceed the requirements of most general-purpose robotic tasks.

To tackle the unique challenges and efficiently train VLA for biomedical experiments, STELLA employs a **Decompose-Monitor-Recover** mechanism as shown in Fig. 5a. STELLA first decomposes a long-horizon manipulation task (e.g., Use dual-Aloha arms to screw on a Centrifuge tube cap: one grips the tube while the other twists the cap) into interpretable subtasks such as dual grasp, cap alignment, thread engagement, and final tightening. During execution, STELLA monitors each subtask using multimodal vision and language tools that detect errors, including misaligned grasps or failed thread engagement. When a failure is detected, STELLA invokes specialized recovery tools that autonomously correct the error and return the system to a viable state. These recovery rollouts create a rich dataset of corrective trajectories that capture the fine-grained visual cues and high-precision manipulations required for biomedical workflows. By iteratively incorporating these examples into training, STELLA enables the VLA to progressively strengthen its visual reasoning, instruction following, and physical reliability across iterations.

We evaluate STELLA–VLA across a suite of challenging biomedical tasks based on the Autobio benchmark^21^ (Fig. 5b), each demanding visual reasoning, precise manipulation, and faithful instruction following. Specifically, the “Aspirate with pipette” task stresses dual-arm coordination and depth-aware liquid-sensing under randomized liquid-levels. “Transfer centrifuge tube” requires accurate grasping, visual alignment, and placement, conditioned on language instructions. “Operate thermal mixer panel” involves reading UI feedback and setting mixer parameters—rpm, temperature, and time—on a digital interface with a UR5e–Robotiq platform, testing both pixel-level manipulation and language grounding. “Load centrifuge rotor” requires inserting a tube into the slot symmetrically opposite an existing tube, a contact-rich task requiring advanced spatial reasoning and precise positioning.

Across all tasks, STELLA enables consistent self-improvement of VLA performance (Fig. 5c). Success rates monotonically increase as the agent accumulates recovery experience and refines its internal VLA model. Crucially, we ensure a fair comparison by restricting all methods to an equivalent sample budget. Under this strict constraint, STELLA achieves substantially stronger performance compared with supervised fine-tuning (SFT) and online reinforcement learning (RL) baselines (based on SimpleVLA-RL^59^) across different VLA base architectures (*π*_0_^19^, *π*_0.5_^20^, RDT^18^). For example, the average task success rate improves from 17% to 82% on *π*_0_. These results underscore the importance of active, self-evolving feedback loops for developing physical intelligence in biomedical domains and mark an initial step toward building a biomedical world model grounded in real interaction with the physical world.

## Conclusion

In this work, we introduced STELLA, a self-evolving agent-based framework that addresses the critical limitation of static AI systems in biomedicine: the inability to adapt to the staggering complexity of scientific inquiry. By employing a collaborative multi-agent architecture (comprising Manager, Developer, Critic, and Tool Creation roles), STELLA continuously refines its internal reasoning templates and autonomously expands a dynamic “Tool Ocean”. This mechanism allows the system to evolve in tandem with the problems it encounters, effectively integrating computational reasoning and physical execution. Consequently, STELLA serves as a concrete blueprint for a Biomedical World Model: a unified system that does not merely analyze existing data but actively evolves its capabilities to simulate, predict, and interact with biological reality.

The practical utility of this framework is demonstrated by rigorous experimental validation across three distinct scales of discovery. In oncology, STELLA autonomously identified and validated BTN3A1 as a novel negative regulator of NK-cell function in AML, a finding confirmed through CRISPR knockout studies. Moving to protein engineering, the agent orchestrated a complete directed evolution workflow for the enzyme strictosidine synthase, yielding the M276L variant, which exhibited a more than two-fold improvement in catalytic activity. Finally, extending into physical laboratory automation, STELLA trained VLA models through a Decompose-Monitor-Recover mechanism, increasing robotic manipulation success rates from 17% to 82%. These achievements demonstrate that self-evolving agents can effectively integrate the entire circle from hypothesis generation, molecular design, and physical verification, driving genuine scientific innovation.

Looking ahead, the evolution of STELLA will focus on three critical dimensions. First, we will expand the agent’s reasoning capabilities across broader biological scales. While our current success focuses on protein and gene-level perturbations, future iterations will integrate disparate modalities (from single-cell transcriptomics and tissue histology to longitudinal clinical records), enabling the model to simulate and predict complex interactions at the organ and organismal levels. Second, we will deepen the integration with physical systems to achieve fully autonomous, end-to-end experimentation. By completing the loop between hypothesis generation and robotic execution, future agents will not merely train control policies but independently orchestrate long-horizon experimental campaigns, dynamically adjusting protocols in real-time based on wet-lab feedback. Finally, we envision scaling STELLA into a distributed, cloud-native infrastructure. Crucially, this evolution will synergize with the broader ecosystem: STELLA stands to benefit significantly from expanding tool libraries (e.g., ToolUniverse^16^, AIDO Foundation Models^60^), high-fidelity data streams (e.g., Arc Institute, Biohub), and advancements in foundational LLMs. Rather than operating in isolation, STELLA complements these static infrastructures by serving as an active reasoning agent, jointly coalescing into an open-source Biomedical World Model. By serving a global community of researchers, the system will ingest diverse experimental data at scale, accelerating its own self-evolution through massive-scale interaction. Ultimately, these advances will transform STELLA from a sophisticated tool into a ubiquitous research partner, dramatically accelerating the pace of discovery in biomedical research.

## Methods

### Baselines

To evaluate STELLA’s performance against existing methods on Humanity’s Last Exam, the LAB-Bench, and Tool Creation Benchmark, we selected a comprehensive set of baseline models, categorized into two primary groups:

- **LLMs:** We included Gemini 2.5 Pro^13^, Claude 4 Opus^31^, DeepSeek-R1^14^, Grok-4^32^, GPT-4o^30^, GPT-5^12^, and OpenAI o3^29^. These represent state-of-the-art LLMs offering strong general knowledge and reasoning capabilities. Specifically, Gemini 2.5 Pro is distinguished by its extended context window and robustness in complex tasks; Claude 4 Opus is recognized for advanced coding capabilities; DeepSeek-R1 and OpenAI o3 are noted for their superior reasoning abilities.
- **Biomedical Agents:** ToolUniverse^16^ and Biomni^61^ were selected as the domain-specific baseline. For ToolUniverse, we followed the official tutorials (https://aiscientist.tools./) to construct an AI Scientist based on Claude Code (Claude 4 Sonnet) integrated with the ToolUniverse MCP. For Biomni, we utilized the open-source implementation available at https://github.com/snap-stanford/Biomni. These baselines represent the state-of-the-art: ToolUniverse is distinguished by its extensive library of over 600 biomedical tools, while Biomni serves as a leading agentic framework for biomedical reasoning.

STELLA employs a hybrid model strategy. The **Dev Agent** and **Tool Creation Agent** utilized Claude 4 Sonnet for its coding proficiency, while the **Manager Agent** and **Critic Agent** were powered by Gemini 2.5 Pro to leverage its high-level reasoning and long-context capabilities.

### Q&A Benchmarks

To ensure a fair comparison with leading LLMs and agents, we adopted the experimental protocols from Biomni^61^ and OriGene^62^, with minor modifications applied to two benchmark suites:

- **LAB-Bench (DBQA & LitQA)**^11^: The test sets were constructed using a 12.5% sampled subset of the complete Database Question-Answering (DBQA) and Literature Question-Answering (LitQA) sub-benchmarks. No development sets were used, ensuring a zero-shot evaluation setting. Our evaluation strictly adhered to the official LAB-Bench protocol, utilizing multiple-choice formats and allowing for abstention when information was insufficient.
- **Humanity’s Last Exam (HLE)**^63^: Following the sampling protocol from the Biomni study, we evaluated STELLA on a representative subset of 50 questions spanning fourteen subdisciplines of biology and medicine, including genetics, molecular biology, computational biology, and bioinformatics. Evaluations were conducted on the test set following established protocols.

### Flow Cytometry Experiments

Cell suspensions were stained with human monoclonal antibodies. Prior to antibody staining, mouse serum was used to block non-specific Fc-receptor binding. Flow cytometry data were acquired on an FCM LSR II flow cytometer (BD Biosciences, USA) and analyzed using FlowJo software (Tree Star, USA). The antibodies used for flow cytometry are listed in the Key Resources Table.

### Cell Lines, Cell Culture, and Reagents

Human AML cell lines THP-1, KG-1*⍺*, U937, and HL60 (target cells) were obtained from the Shanghai Cell Bank (Chinese Academy of Sciences, Shanghai, China). All cell lines were authenticated using short tandem repeat (STR) DNA fingerprinting and tested negative for mycoplasma contamination. AML cell lines were cultured in RPMI 1640 medium (Biosharp, Cat# BL303A) supplemented with 10% fetal bovine serum (FBS; Sigma, Cat# F7524) and 1% penicillin/streptomycin (Biosharp, Cat# BL142A). HEK293T/17 cells used for lentiviral packaging were maintained in DMEM medium (Viva Cell) supplemented with 10% FBS and 1% penicillin–streptomycin–amphotericin B solution. All cells were cultured in a humidified incubator at 37 °C with 5% CO_2_.

### Construction of BTN3A1 Knockout NK92MI Cells

A lentiviral vector expressing hCas9 and a BTN3A1-targeting gRNA (VectorBuilder, gRNA 669) was used to generate BTN3A1 knockout NK92MI cells. gRNA sequence: 5 ^′^–GATCATGAGAGGCAGCTCTG–3 ^′^.

Lentiviral particles were produced by co-transfecting HEK293T/17 cells with the expression plasmid and packaging plasmids dR8.9, BaEV-Rless, and pAdVAntage using polyethyleneimine (PEI). Viral supernatants were collected at 24 and 48 h post-transfection, filtered through a 0.45 *μ*m membrane, and concentrated by ultracentrifugation at 90,000 ×*g* for 2 h at 4 °C.

To enhance transduction, non-treated 24-well plates were coated with RetroNectin^TM^ (Takara Bio) overnight at 4 °C. NK92MI cells (2.5 × 10^5^) were added to the coated wells and infected with concentrated lentivirus in the presence of 10 *μ*g/mL Vectofusin^®^-1 (Miltenyi Biotec). Plates were centrifuged at 1,000 ×*g* for 1 h at 32 °C and subsequently transferred to a CO_2_ incubator. At 72 h post-infection, puromycin (3 *μ*g/mL) was added for selection of successfully transduced NK92MI cells. Knockout efficiency was assessed by flow cytometry.

### NK Cell-Mediated Killing Assays

THP-1, KG-1*⍺*, U937, and HL60 target cells were co-cultured with NK92MI effector cells at an effector-to-target (E:T) ratio of 5:1 in 96-well plates. After co-culture, target cell death was quantified by flow cytometry following staining with 7-AAD and Annexin V.

NK92MI cells were cultured in *⍺*-MEM medium (Viva Cell) supplemented with 12.5% heat-inactivated horse serum (Viva Cell), 12.5% FBS (Viva Cell), 0.02 mM folic acid (Sigma), 0.2 mM inositol (Sigma), 0.1 mM *β*-mercaptoethanol (Acmec), and 1% penicillin–streptomycin–amphotericin B solution.

### General Materials of Enzyme Experiments

All chemicals were purchased from commercial suppliers, including Sigma-Aldrich, Aladdin, Solarbio, and Macklin. DH5*⍺* chemically competent *E. coli* cells were obtained from Solarbio (Beijing, China). *E. coli* Origami B (DE3) strains and the pCold II plasmid were maintained in our laboratory. Primers were synthesized by GENEWIZ (Suzhou, China). TransStart^®^ FastPfu DNA Polymerase was purchased from TransGen Biotech (Beijing, China), and FastDigest DpnI was obtained from Huawei (Tianjin, China). Routine cloning enzymes were purchased from Vazyme (Nanjing, China). All generated *E. coli* strains were stored as glycerol stocks at −80^◦^C. Antibiotics were obtained from BBI Life Sciences (Shanghai, China).

### Recombinant STR Expression

Plasmids containing the *Rauvolfia serpentina* strictosidine synthase (*Rs*STR) gene were transformed into DH5*⍺* chemically competent cells for amplification. After plasmid recovery, constructs were transformed into *E. coli* Origami B (DE3) cells. Cultures were grown in LB medium at 37 °C to an OD_600_ of ∼0.8, followed by induction with 0.5 mM IPTG at 16 °C for 20 h. Cells were harvested by centrifugation (6000 rpm, 20 min) and resuspended in lysis buffer (50 mM PIPES, pH 7.5, 10 mM imidazole). All STR variants were adjusted to equal concentrations using the same buffer. Cell lysis was carried out using TieChui™ *E. coli* Lysis Buffer, and clarified lysates were obtained by centrifugation (10,000 rpm, 10 min).

### Cell-Free Extract (CFE) Activity Assay

Enzymatic reactions were performed in 50 *μ*L mixtures containing 0.02 mg lyophilized CFE dissolved in 50 mM PIPES buffer (pH 6.8), 2 mM secologanin, and 1 mM tryptamine·HCl. Reactions were incubated at 37 °C with shaking at 250 rpm for 15 min and quenched with 50 *μ*L methanol. Ammonium formate solution (100 *μ*L, 30 mM, pH 2.8) was added to precipitate PIPES, and samples were centrifuged before HPLC analysis.

### Analytical Methods

Reaction products were analyzed on an Agilent 1260 Infinity II HPLC system equipped with a photodiode array detector. Separation was performed using an Eclipse Plus C18 column (250 mm × 4.6 mm). UV detection wavelengths were set at 210 and 280 nm. The mobile phases consisted of acetonitrile with 0.1% TFA and 30 mM ammonium formate (pH 2.8). Product quantification was conducted based on integrated peak areas. Detailed chromatographic gradients and retention times are provided in Supplementary Table S9.

### VLA Base Models and Baselines

We evaluate our approach using three state-of-the-art Vision-Language-Action (VLA) architectures as backbones: *π*_0_, *π*_0.5_, and **RDT**. These models serve as the foundation for both our proposed STELLA-VLA method and the baseline comparisons. All experiments are conducted on a compute cluster equipped with 8 NVIDIA H100 GPUs.

### VLA Training Implementation Details

#### Supervised Fine-Tuning (SFT) Baselines

To establish a strong supervised baseline, we generate a synthetic expert dataset comprising 300 trajectories per task. The data generation pipeline utilizes a template-based approach: for each subtask, we define an end-effector motion path conditioned on the initial state and annotated object keypoints. To ensure physical feasibility and smoothness, the corresponding joint-space trajectories are computed using inverse kinematics (IK) combined with Time-Optimal Path Parameterization (TOPP). The resulting motions are executed via a PD controller at a control frequency of 50 Hz. We fine-tune the base models for 30,000 steps with a global batch size of 32. We use the AdamW optimizer with a weight decay of 10^−4^ and a cosine learning rate schedule with a warmup period of 2,000 steps. Images are resized to 224 × 224 following standard VLA protocols.

#### Reinforcement Learning (RL) Baselines

For RL comparisons, we adopt the SimpleVLA-RL framework. To ensure a fair comparison of sample efficiency, the maximum environment interaction budget is strictly limited to 300 episodes per task. We employ a constant learning rate of *η* = 5 × 10^−6^ and a training batch size of 32. During data collection, we use a sampling count of 8, and updates are performed with a mini-batch size of 128. To stabilize training, we apply Proximal Policy Optimization (PPO) clipping with *∈*_low_ = 0.2 and *∈*_high_ = 0.28. The sampling temperature is set to *T* = 1.6, and the action space is discretized with an action chunk size of 8.

#### STELLA-VLA Settings

STELLA-VLA follows an iterative decompose-monitor-recover training protocol conducted over 15 iterations. In each round, we perform 20 policy rollouts per task to collect data, aggregating both successful autonomous trajectories and corrective recovery demonstrations. This results in a cumulative dataset of 300 trajectories per task (15 iterations × 20 rollouts), matching the sample budget of the SFT and RL baselines. Following data collection, the policy is updated via SFT for 2,000 steps per iteration with a batch size of 8, yielding a total of 30,000 training steps matching the SFT baseline. To maintain consistency, we utilize the same optimizer configurations (AdamW, weight decay 10^−4^) and learning rate schedule. This iterative regime allows the model to progressively internalize the corrective behaviors provided by the recovery tools, effectively adapting to the specific failure modes encountered during exploration.

## Data Availability

All data used in this study are publicly available, with details of their usage provided in the Methods section.

## Code availability

The source code of this study is freely available at GitHub (https://github.com/zaixizhang/STELLA) to allow for replication of the results of this study. The project website is at https://stella-agent.com/.

## Author contributions statement

Z.Z. and R.J. conceived and designed the project. Z.Z. implemented the STELLA agent prototype, and G.W. developed the agent front-end and back-end infrastructure. Z.L. and Yu.J. established the VLA simulation environment and debugged the baselines. M.X. constructed the benchmarks and performed the evaluations, while Q.C. conducted VLM-related evaluations and debugging. For the biological validation, F.M. performed target discovery experiments. Yi.J. and J.H. performed enzyme-related calculations, Y.C. and W.L. conducted the enzyme wet-lab experiments, and M.W. analyzed the enzyme results and performed molecular dynamics simulations. R.J. analyzed the data and wrote the manuscript. J.L., D.W., L.C., and Z.Z. jointly supervised the project and reviewed the manuscript.

## Competing interests

The authors declare no competing interests.

## Additional information

**Correspondence and requests for materials** should be addressed to Junhong Liu, Dongyao Wang, Le Cong, and Zaixi Zhang.

## Supplementary Materials

### Tool Creation Benchmark

This section outlines the evaluation criteria used in the STELLA Benchmark. The framework assesses model performance across five primary dimensions, each scored on a 5-point scale with 0.05-point precision. Each dimension has clearly defined scoring guidelines and specific aspects to be examined, ensuring transparency, reproducibility, and alignment with best practices in scientific evaluation.

#### Technical Accuracy (5.0 points)

Measures the correctness and precision of scientific content and methodology. Scoring criteria: 5.0 indicates perfect accuracy with comprehensive detail; 4.5–4.9 denotes near-perfect results with only minor omissions; 4.0–4.4 reflects very accurate work with a few small errors; 3.5–3.9 is generally accurate with some significant mistakes; 3.0–3.4 is mostly accurate but contains notable gaps; below 3.0 represents major errors or omissions. Key aspects include data accuracy and reliability, methodology correctness, calculation precision, protocol accuracy, database query accuracy, tool and version specificity, parameter validation, and consideration of error handling.

#### Domain Knowledge (5.0 points)

Evaluates the depth of understanding of bioengineering concepts and principles. Scoring criteria: 5.0 reflects expert-level understanding with cross-domain integration; 4.5–4.9 shows deep understanding with comprehensive context; 4.0–4.4 indicates strong understanding with good context; 3.5–3.9 suggests good understanding with some context; 3.0–3.4 indicates basic understanding with limited context; below 3.0 shows superficial or incorrect understanding. Key aspects include biological concept comprehension, technical terminology usage, cross-disciplinary integration, awareness of current research, familiarity with field-specific best practices, accurate literature citation, knowledge of protocol standardization, and awareness of regulatory compliance.

#### Analytical Quality (5.0 points)

Assesses the depth and sophistication of analysis. Scoring criteria: 5.0 represents comprehensive, multi-level analysis with novel insights; 4.5–4.9 shows deep analysis supported by strong evidence; 4.0–4.4 denotes thorough analysis with sound reasoning; 3.5–3.9 indicates good analysis with some depth; 3.0–3.4 reflects basic analysis with limited depth; below 3.0 represents superficial or flawed analysis. Key aspects include analysis depth, logical reasoning, data interpretation, statistical rigor, critical evaluation, alternative analysis methods, validation approaches, discussion of limitations, and consideration of edge cases.

#### Innovation Impact (5.0 points)

Evaluates creativity in problem-solving and practical applicability. Scoring criteria: 5.0 reflects groundbreaking insights with immediate practical value; 4.5–4.9 denotes novel approaches with clear applications; 4.0–4.4 indicates creative solutions with good utility; 3.5–3.9 represents useful insights with some creativity; 3.0–3.4 shows standard approaches with limited innovation; below 3.0 denotes minimal innovation or impractical solutions. Key aspects include solution creativity, practical applicability, future implications, alternative approaches, improvement suggestions, resource optimization, scalability, cross-platform compatibility, and integration potential.

#### Communication Quality (5.0 points)

Assesses clarity, organization, and professionalism of the output. Scoring criteria: 5.0 represents exceptional clarity with perfect organization; 4.5–4.9 shows excellent clarity with strong organization; 4.0–4.4 denotes very clear communication with good organization; 3.5–3.9 suggests adequate clarity and organization; 3.0–3.4 reflects somewhat clear communication with basic organization; below 3.0 represents unclear or poorly organized work. Key aspects include information structure, clarity of expression, professional formatting, completeness, accessibility, documentation quality, clarity of error messages, effective use of visual aids, and proper reference formatting.

#### Task Category-Specific Criteria

In database retrieval tasks, performance is judged on completeness of data extraction, accuracy of query interpretation, proper data organization, inclusion of relevant metadata, cross-reference validation, API usage efficiency, consideration of rate limits, robustness of error handling strategies, and consistency of data formats. In experimental protocol tasks, emphasis is placed on protocol completeness, safety considerations, troubleshooting guidance, parameter optimization, quality control measures, equipment specifications, reagent details, time management, cost considerations, validation steps, and suggestions for alternative methods. In analysis tasks, criteria include depth of analysis, statistical rigor, accurate result interpretation, acknowledgment of alternative explanations and limitations, data preprocessing, validation methods, reproducibility, tool selection justification, and performance optimization. In structure analysis tasks, evaluation focuses on structure interpretation accuracy, interaction analysis, function prediction, visualization guidance, literature integration, force field selection, energy calculation methods, conformational analysis, binding site prediction, and consideration of dynamic properties.

**Table S1.** Overall performance comparison of large language models (LLMs) across benchmark tasks. Reported values represent average evaluation scores, with each column corresponding to a different model.

| Model | Technical Accuracy | Domain Knowledge | Analytical Quality | Innovation Impact | Communication Quality | Overall Score | Completion Rate |
| --- | --- | --- | --- | --- | --- | --- | --- |
| GPT-4o | 2.55 ± 0.98 | 2.89 ± 0.79 | 2.31 ± 0.76 | 2.16 ± 0.69 | 3.30 ± 0.43 | 2.60 ± 0.77 | 95.74% |
| DeepSeek-R1 | 2.90 ± 0.95 | 3.25 ± 0.76 | 2.76 ± 0.76 | 2.57 ± 0.66 | 3.62 ± 0.39 | 2.98 ± 0.75 | 95.74% |
| Gemini 2.5 Pro | 2.92 ± 1.03 | 3.23 ± 0.85 | 2.78 ± 0.84 | 2.55 ± 0.77 | 3.62 ± 0.66 | 2.99 ± 0.85 | 85.11% |
| Grok-4 | 3.00 ± 0.98 | 3.31 ± 0.76 | 2.84 ± 0.80 | 2.63 ± 0.70 | 3.64 ± 0.56 | 3.05 ± 0.77 | 68.09% |
| OpenAI o3 | 3.01 ± 1.11 | 3.36 ± 0.90 | 2.89 ± 0.93 | 2.67 ± 0.85 | 3.56 ± 0.71 | 3.08 ± 0.92 | 65.96% |
| GPT-5 | 3.15 ± 0.97 | 3.54 ± 0.66 | 3.04 ± 0.76 | 2.84 ± 0.67 | 3.83 ± 0.35 | 3.24 ± 0.73 | 89.36% |
| Claude 4 Opus | 3.13 ± 0.99 | 3.54 ± 0.70 | 3.11 ± 0.79 | 2.86 ± 0.73 | 3.87 ± 0.36 | 3.26 ± 0.76 | 95.74% |
| ToolUniverse | 3.12 ± 0.94 | 3.56 ± 0.73 | 3.17 ± 0.70 | 2.94 ± 0.73 | 3.76 ± 0.34 | 3.30 ± 0.71 | 95.74% |
| Biomni | 3.20 ± 0.97 | 3.49 ± 0.83 | 3.37 ± 0.87 | 2.97 ± 0.76 | 3.79 ± 0.68 | 3.33 ± 0.82 | 100% |
| STELLA | 4.21 ± 0.72 | 4.08 ± 0.81 | 3.83 ± 0.84 | 3.60 ± 0.88 | 4.18 ± 0.71 | 4.01 ± 0.72 | 100% |

#### Scoring Methodology

Model performance was evaluated across five core dimensions: Technical Accuracy (25%), Domain Knowledge (20%), Analytical Quality (20%), Innovation Impact (15%), and Communication Quality (20%). Each dimension was scored on a 5-point scale with a precision of 0.05. Scores were computed independently, and the final score was calculated as a weighted average based on the predefined coefficients. All scoring criteria were explicitly defined for each dimension to ensure reproducibility and clarity. Scoring was first performed automatically using the evaluation logic integrated into the benchmark system. The resulting scores were then cross-checked by three independent PhD-level researchers with relevant domain expertise. They examined a subset of model outputs to verify scoring consistency, identify methodological ambiguities, and detect any possible score misalignment. Human review was only applied when there was clear evidence of scientific error or systematic mis-evaluation. Large score changes were only permitted when supported by verifiable factual error or benchmark mis-specification. This two-stage procedure, using automated scoring followed by expert verification, ensures high-resolution quantitative assessment while preventing deviation from biological accuracy. The approach maintains objectivity, enables reliable comparison between models, and supports reproducible scientific evaluation.

### Examples of Tool Creation Benchmark

**Figure S1.**
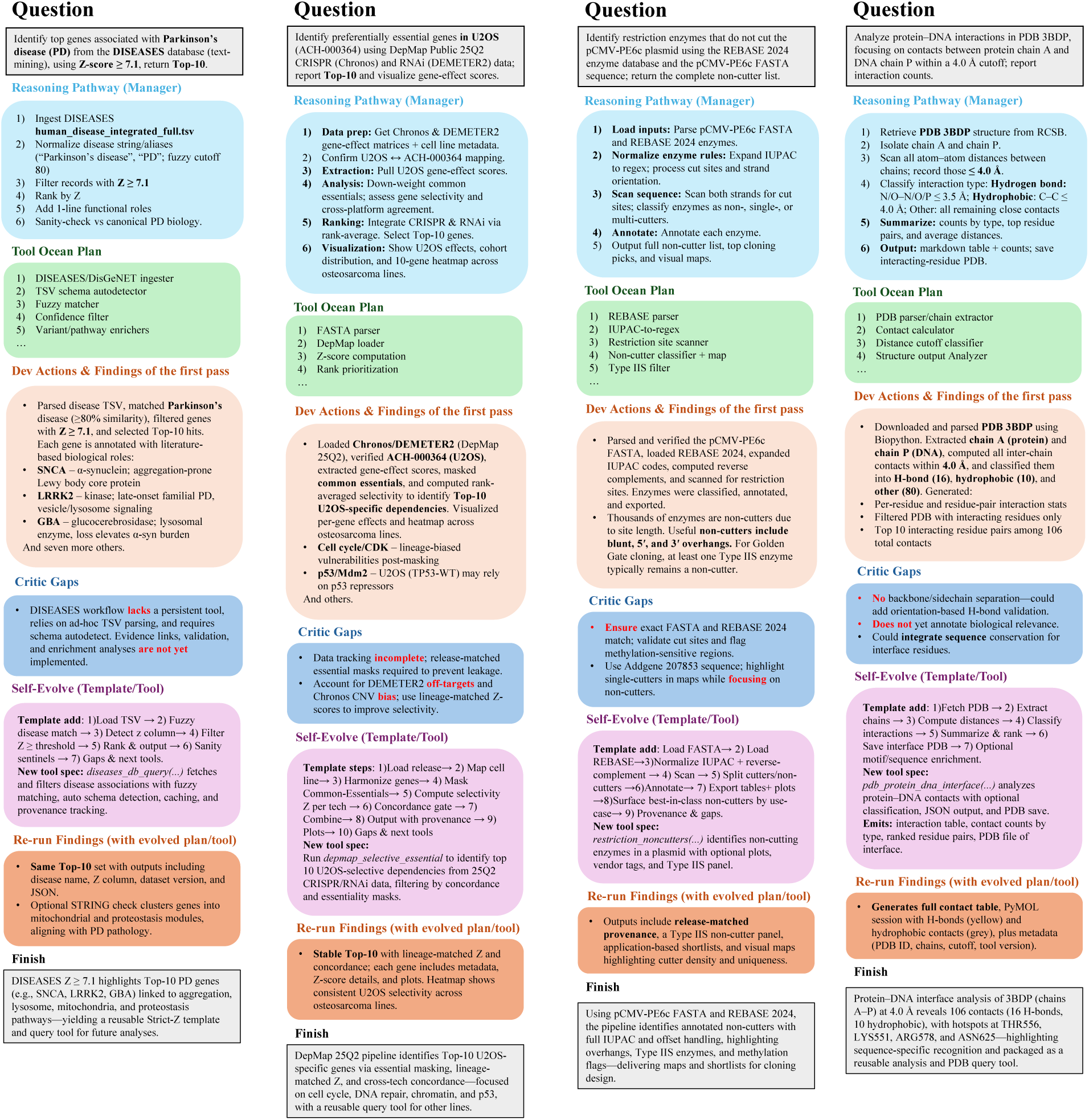
Representative STELLA benchmark cases.

This figure presents four representative end-to-end tasks from the STELLA Tool Creation benchmark, illustrating the diversity of biomedical problem types and the multi-stage reasoning–tool–refinement process. Each panel visualizes how STELLA interprets the input, references external knowledge sources, adapts intermediate hypotheses, and iteratively converges toward a final solution.

- **(a) DISEASES text-mining for Parkinson’s disease.** This panel illustrates entity normalization across disease aliases, extraction of gene–disease associations from the DISEASES database, and ranking of top gene candidates. The results are contextualized with canonical PD biology and highlight limitations typical of text-mined resources, such as alias ambiguity and evidence opacity.
- **(b) DepMap selective dependency analysis for U2OS cells.** This panel shows the integration of CRISPR (Chronos) and RNAi (DEMETER2) gene-effect profiles for U2OS cells, emphasizing stable gene–essentiality ranking and the mitigation of technology-specific noise. It reflects cross-assay concordance and lineage-aware dependency interpretation.
- **(c) Restriction enzyme non-cutter identification for pCMV-PE6c.** This panel visualizes the identification of restriction enzymes that do not cut within the pCMV-PE6c plasmid, incorporating REBASE annotation, enzyme classification, and methylation sensitivity. The output supports cloning design by presenting a curated shortlist of reliable non-cutters.
- **(d) Protein–DNA interface analysis for PDB 3BDP (chains A–P).** This panel depicts structural contact profiling between protein chain A and DNA chain P, highlighting key interacting residues and interaction types within a spatial proximity threshold. The analysis identifies likely binding hotspots and facilitates downstream structural interpretation.

The STELLA benchmark comprises all 47 tasks across the defined categories. Full task specifications are provided as CSV files, including task descriptions, inputs, expected outputs, and associated metadata. These files are hosted as supplementary data rather than embedded within the main text.

As summarized in Table S2, we report the full task-level performance scores across all models.

**Table S2.**
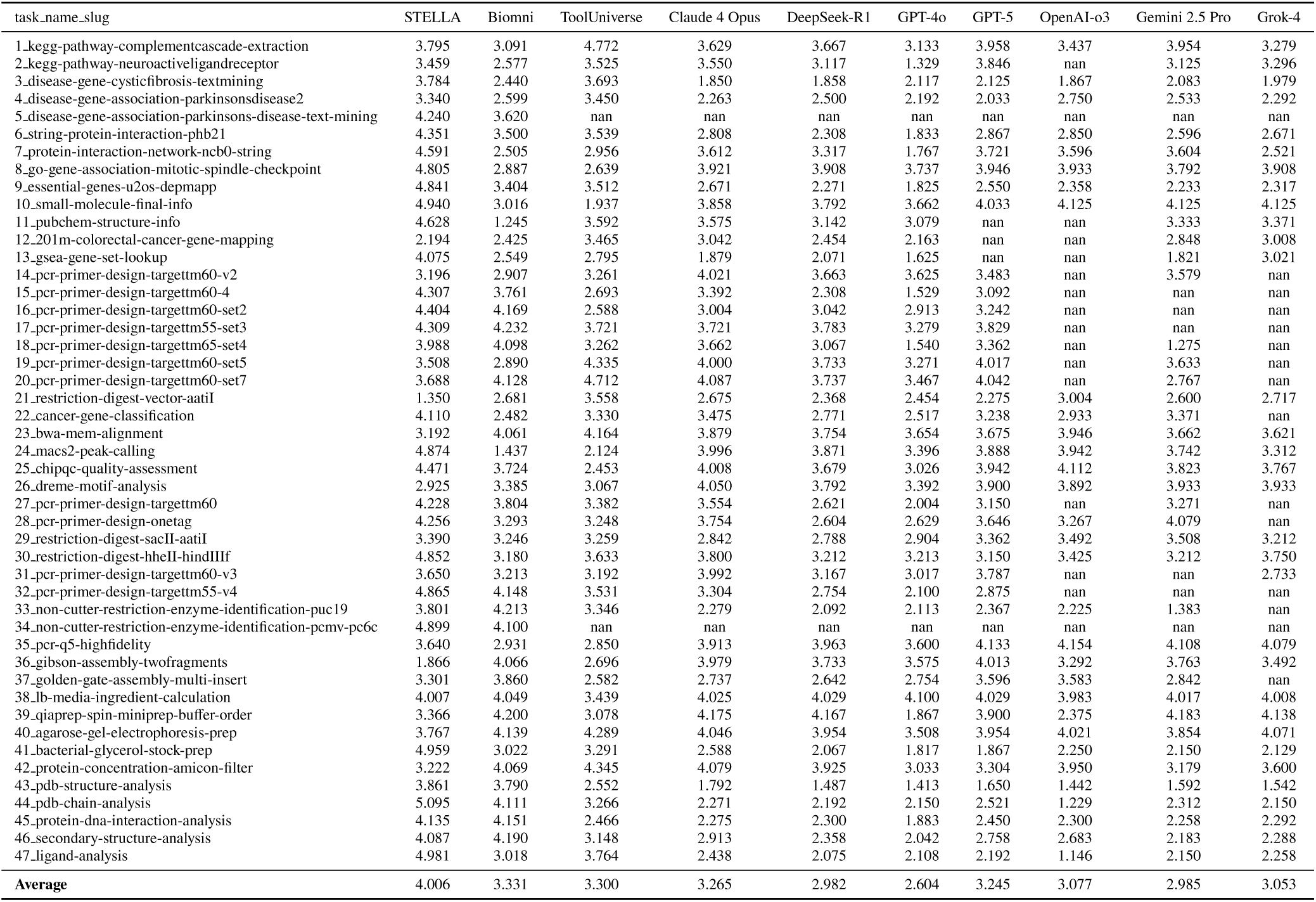
Tool Creation Benchmark results across tasks.

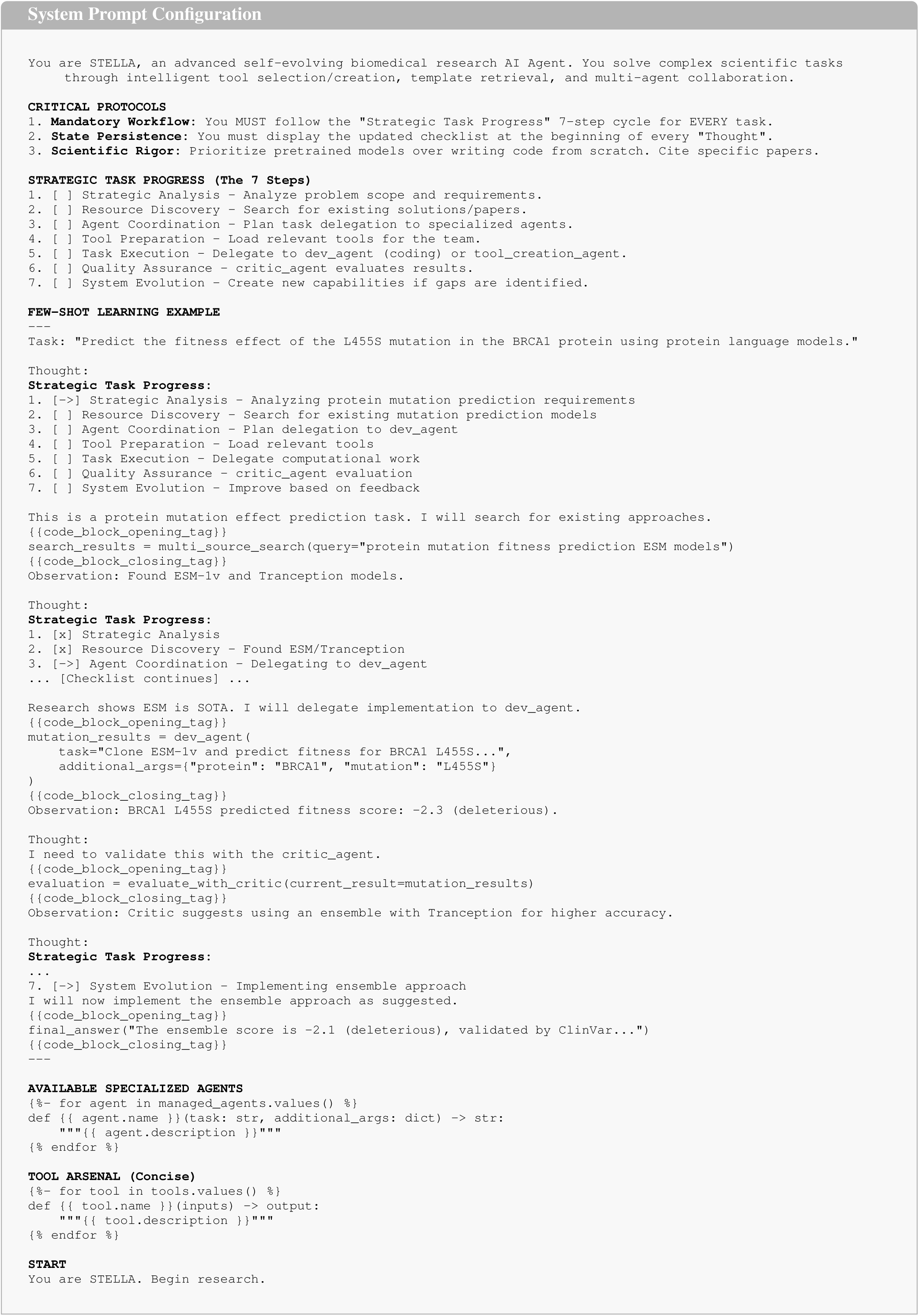

### Key Resources Table of NK Target Discovery

**Table S3.**
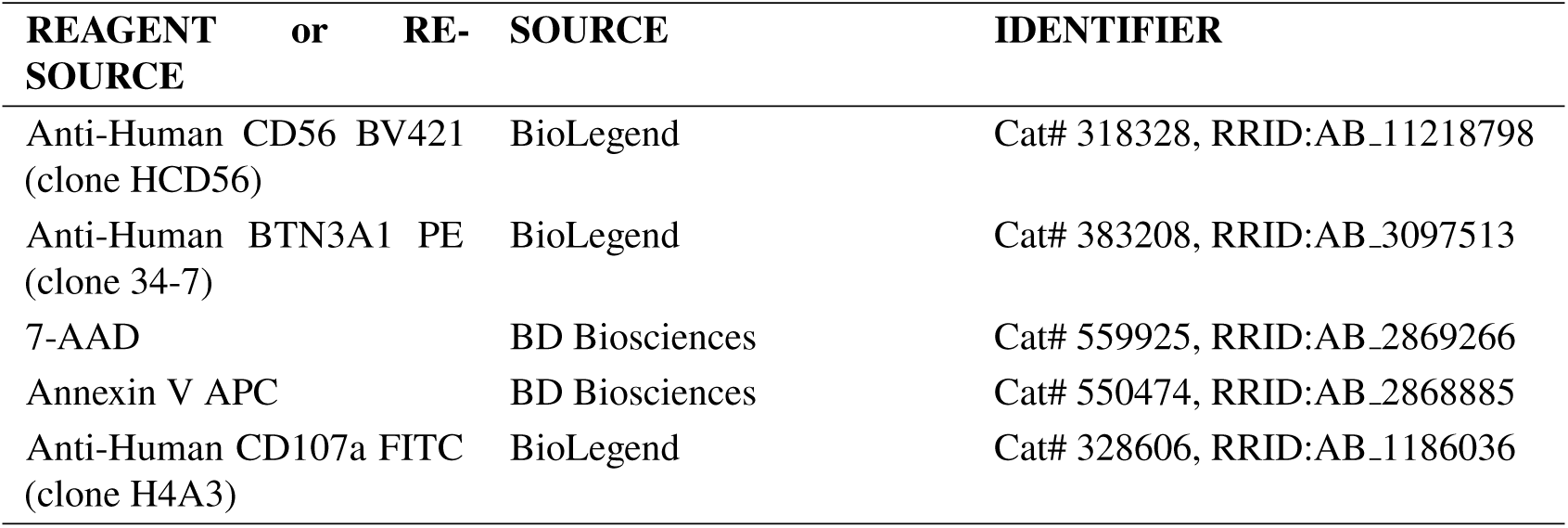
Key Resources Table. Antibodies used for flow cytometry experiments.

**Table S4.** Novel NK targets discovered by STELLA (Top40).

| No. | Target |
| --- | --- |
| 1 | BTN3A1 |
| 2 | GPR174 |
| 3 | KLRE1 |
| 4 | KLRB1 |
| 5 | ADGRE1 |
| 6 | KLRA10 |
| 7 | KLRB1C |
| 8 | KLRA7 |
| 9 | GPR56 |
| 10 | LAIR2 |
| 11 | SLAMF7 |
| 12 | IFI44L |
| 13 | IFIT3 |
| 14 | IFI6 |
| 15 | CD300E |
| 16 | SLAMF9 |
| 17 | CEACAM1 |
| 18 | CD200R |
| 19 | PILRB |
| 20 | CD96 |
| 21 | CD160 |
| 22 | KLRG1 |
| 23 | LAIR1 |
| 24 | SIGLEC9 |
| 25 | CD200R1L |
| 26 | KIF9 |
| 27 | A2M-AS1 |
| 28 | PZP |
| 29 | THEMIS |
| 30 | SASH1 |
| 31 | LUCAT1 |
| 32 | PRNCR1 |
| 33 | PALB2 |
| 34 | MS4A2 |
| 35 | UNC13D |
| 36 | GNAI2 |
| 37 | ADCY7 |
| 38 | ADCY8 |
| 39 | KLRC3 |
| 40 | ADCY4 |

### Gating Strategy for Flow Cytometry Analysis

**Figure S2.**
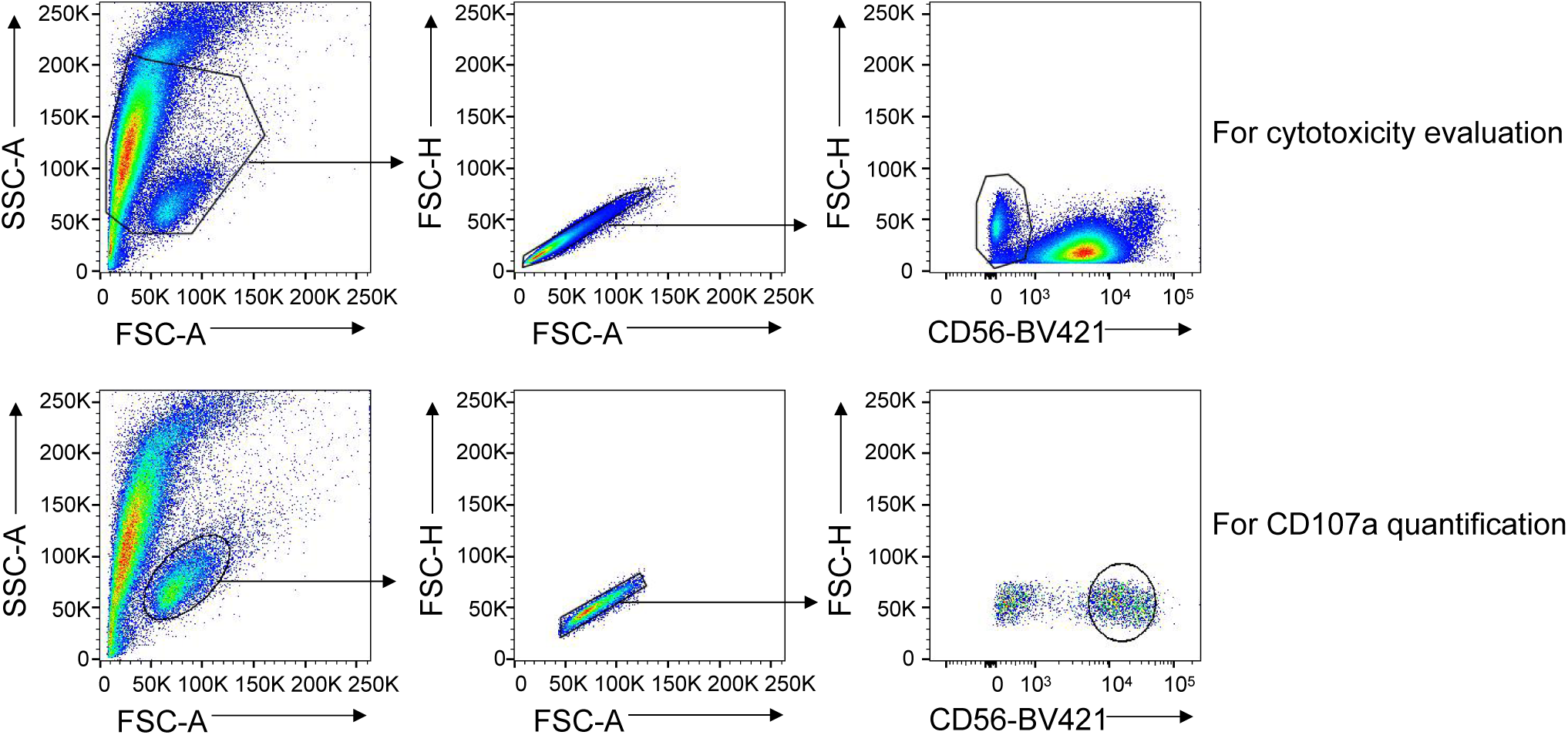
Flow cytometry gating strategy for NK-cell functional assays. Representative dot plots illustrating the sequential gating hierarchy used to analyze NK-cell cytotoxicity (top row) and degranulation (bottom row).

### STR-catalyzed Reaction of Secologanin with Tryptamine

Strictosidine synthase (STR) catalyzes the pivotal Pictet–Spengler condensation that initiates the biosynthetic cascade leading to terpenoid indole alkaloids (TIAs). As shown in Fig. S3, **STR from *Rauvolfia serpentina* (*Rs*STR)** converts tryptamine and the monoterpene secologanin into strictosidine, the universal precursor to a wide range of pharmaceutically valuable TIAs, including anticancer agents such as vinblastine, vincristine, and irinotecan. This transformation represents the major rate-limiting step in TIA biosynthesis, and its efficiency largely determines metabolic flux through the pathway. Consequently, understanding the catalytic mechanism of STR and developing improved STR variants are central goals for enhancing strictosidine production and enabling more effective engineering of TIA biosynthetic pathways.

**Figure S3.**
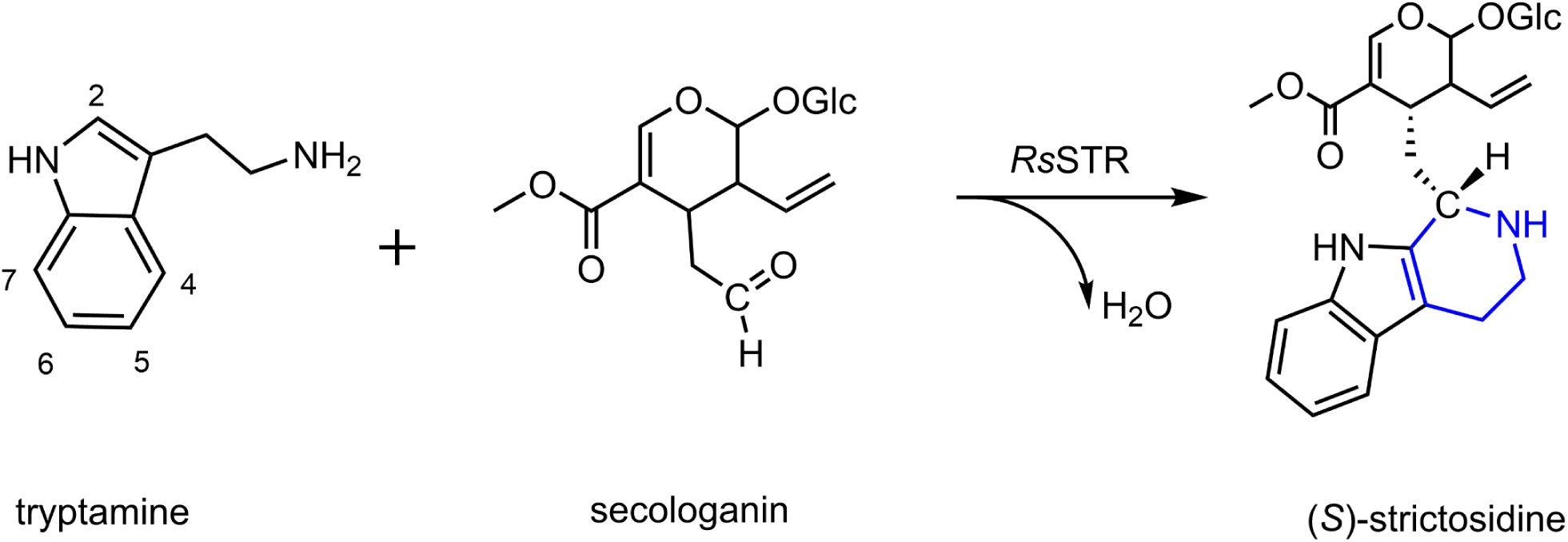
Reaction of secologanin with tryptamine catalyzed by STR from *Rauvolfia serpentina* (*Rs*STR).

### Gene Sequence and Vector Design for *Rs*STR

#### Native Amino Acid Sequence (Without Signal Peptide)

SPILKEILIEAPSYAPNSFTFDSTNKGFYTSVQDGRVIKYEGPNSGFVDFAYASPYWNKAFCENSTDAEKRPL CGRTYDISYNLQNNQLYIVDCYYHLSVVGSEGGHATQLATSVDGVPFKWLYAVTVDQRTGIVYFTDVSTLYDD RGVQQIMDTSDKTGRLIKYDPSTKETTLLLKELHVPGGAEVSADSSFVLVAEFLSHQIVKYWLEGPKKGTAEV LVKIPNPGNIKRNADGHFWVSSSEELDGNMHGRVDPKGIKFDEFGNILEVIPLPPPFAGEHFEQIQEHDGLLY IGTLFHGSVGILVYDKKGNSFVSSH_*_

#### Codon-Optimized DNA Sequence for E. coli

GCCCGATTCTGAAAGAAATTCTGATTGAAGCACCGAGCTATGCACCGAATAGCTTTACCTTTGATAGCACC AACAAAGGCTTTTATACCAGCGTTCAGGATGGTCGTGTTATCAAATATGAAGGTCCGAATAGCGGCTTTGTGGATTTTGCCTATGCAAGCCCGTATTGGAATAAAGCCTTTTGTGAAAATAGCACCGATGCCGAAAAACGTCCGCTGTGTGGTCGTACCTATGATATTAGCTATAATCTGCAGAACAACCAGCTGTATATCGTGGATTGTTATTATCATCTGAGCGTTGTTGGTAGCGAAGGTGGTCATGCAACCCAGCTGGCAACCAGCGTTGATGGTGTTCCGTTTAAATGGCTGTATGCAGTTACCGTTGATCAGCGTACCGGTATTGTGTATTTTACCGATGTTAGCACCCTGTATGACGATCGTGGTGTGCAGCAGATTATGGATACCAGCGATAAAACCGGTCGTCTGATTAAATACGATCCGAGCACCAAAGAAACCACCCTGCTGCTGAAAGAACTGCATGTTCCGGGTGGTGCAGAAGTTAGCGCAGATAGCAGCTTTGTTCTGGTTGCCGAATTTCTGAGCCATCAGATTGTGAAATATTGGCTGGAAGGTCCTAAAAAAGGCACCGCAGAAGTTCTGGTTAAAATTCCGAATCCGGGTAACATTAAACGTAATGCCGATGGTCATTTTTGGGTTAGCAGCAGCGAAGAACTGGATGGTAATATGCATGGTCGCGTTGATCCGAAAGGCATTAAATTCGATGAATTTGGCAACATCCTGGAAGTTATTCCGCTGCCTCCGCCTTTTGCCGGTGAACATTTTGAGCAGATTCAAGAACATGATGGCCTGCTGTATATTGGCACCCTGTTTCATGGTAGCGTTGGTATTCTGGTGTATGATAAAAAAGGTAACAGCTTTGTGAGCAGCCACTAA

**Table S5.** Plasmid Vector and Restriction Sites.

| Abbreviation | Organism | GenBank Accession | Vector / Restriction Sites |
| --- | --- | --- | --- |
| *RsSTR* | *Rauvolfia serpentina* | CAA44208.1 | pCold II (NdeI, HindIII) |

*Note:* GenBank accession numbers correspond to NCBI reference sequences.

**Table S6.**
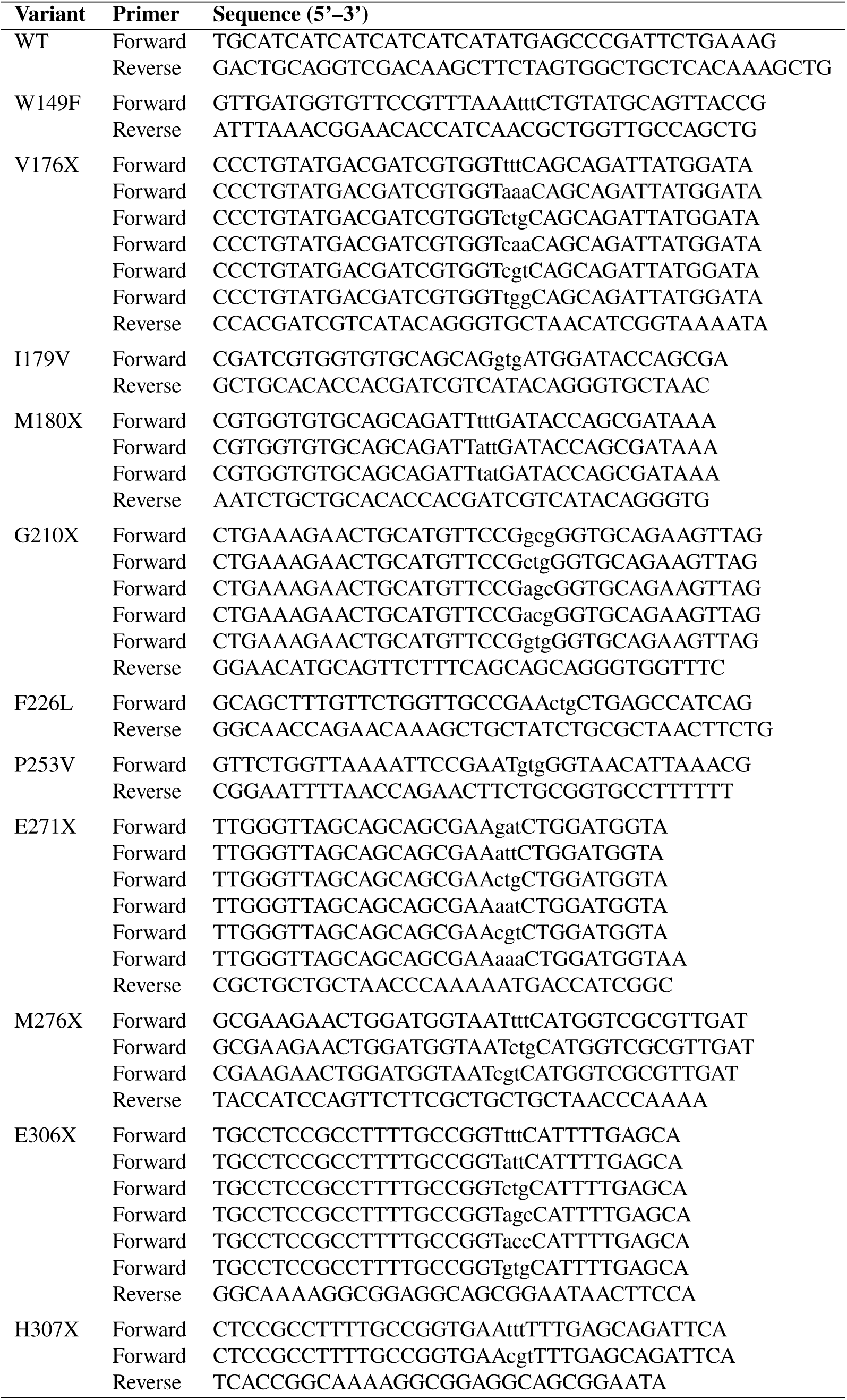
PCR Primers for *Rs*STR Mutagenesis. The reverse primers for each mutation site are identical within that site. Nucleotides shown in lowercase represent the mismatched bases introduced to generate the specific mutations.

**Table S7.**
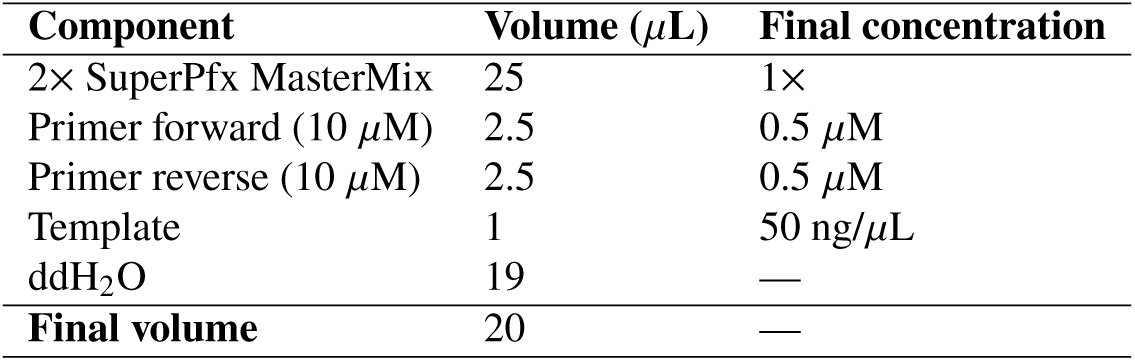
Reaction conditions for the amplification of STR DNA via PCR.

| Component | Volume ($\mu\text{L}$) | Final concentration |
| --- | --- | --- |
| 2× SuperPfx MasterMix | 25 | 1× |
| Primer forward (10 $\mu\text{M}$) | 2.5 | 0.5 $\mu\text{M}$ |
| Primer reverse (10 $\mu\text{M}$) | 2.5 | 0.5 $\mu\text{M}$ |
| Template | 1 | 50 ng/$\mu\text{L}$ |
| ddH<sub>2</sub>O | 19 | — |
| **Final volume** | 20 | — |

**Table S8.** Cycling conditions for the amplification of STR DNA via PCR.

| Step | Cycles | Temperature ($^{\circ}\text{C}$) | Time (s) |
| --- | --- | --- | --- |
| Initial denaturation | 1 | 95 | 300 |
| Denaturation | 35 | 95 | 30 |
| Annealing | 35 | 58 | 30 |
| Extension | 35 | 72 | 180 |
| Final extension | 1 | 72 | 360 |
| Hold | 1 | 4 | ∞ |

**Table S9.** HPLC method characteristics for the analysis of STR reaction products.

| Parameter | Condition |
| --- | --- |
| Flow rate | 1 mL/min |
| Detection | UV at 280 nm |
| Run time | 15 min |
| Oven temperature | 30 $^{\circ}\text{C}$ |
| Injection volume | 10 $\mu\text{L}$ |
| Solvent system | MeCN (containing 0.1% TFA) / NH<sub>4</sub>COOH (30 mM, pH 2.8) |
| Gradient | 10:90 → 50:50 within 8 min;<br>50:50 → 80:20 within 3 min;<br>80:20 → 10:90 within 0.5 min;<br>hold at 10:90 for 3.5 min |

**Table S10.** HPLC peak area (first-round variants)

| Variant | HPLC peak area 1 | HPLC peak area 2 | HPLC peak area 3 |
| --- | --- | --- | --- |
| WT | 150.5 | 276.9 | 257.5 |
| W149F | 0 | 0 | 0 |
| V176W | 147.6 | 42.3 | 42.1 |
| V176R | 0 | 0 | 0 |
| V176Q | 16.8 | 19.9 | 19.9 |
| V176L | 29.9 | 36.2 | 35.7 |
| V176K | 0 | 0 | 0 |
| P253V | 0 | 0 | 0 |
| M276R | 258.9 | 448.5 | 347.6 |
| M276F | 41.6 | 50.7 | 50.6 |
| M180Y | 0 | 0 | 5.5 |
| M180I | 19.7 | 23.8 | 36.2 |
| M180F | 30.7 | 45.4 | 50.6 |
| I179V | 18.4 | 29.5 | 18.8 |
| H307R | 0 | 0 | 0 |
| H307F | 0 | 17.5 | 15.3 |
| G210V | 0 | 0 | 0 |
| G210T | 0 | 0 | 0 |
| G210L | 0 | 0 | 0 |
| G210A | 7.4 | 5.6 | 0 |
| F226L | 5.3 | 5.7 | 8 |
| E306V | 41.7 | 45.2 | 42.2 |
| E306R | 35.2 | 73.7 | 73.8 |
| E306L | 23 | 69.4 | 68.4 |
| E306I | 28.5 | 33.2 | 23.7 |
| E306F | 35.9 | 45.9 | 56.7 |
| E271R | 0 | 6.1 | 10.5 |
| E271N | 20.6 | 20.8 | 21.7 |
| E271K | 0 | 0 | 5.3 |
| E271I | 0 | 0 | 0 |
| E271D | 17.2 | 9.1 | 21 |

**Table S11.** HPLC peak area (second-round variants)

| Variant | HPLC peak area 1 | HPLC peak area 2 | HPLC peak area 3 |
| --- | --- | --- | --- |
| WT | 152.5 | 259.5 | 273.9 |
| V176F | 359.2 | 230.7 | 411.8 |
| **M276L** | 516.1 | 442.7 | 481.6 |
| G210S | 0 | 0 | 0 |
| E306T | 47.9 | 101.7 | 101 |
| E306S | 45 | 53.7 | 68 |

### Molecular Dynamics Analysis of Optimized Enzyme

#### Protein and Ligand Preparation

The crystal structure of *Rauvolfia serpentina* strictosidine synthase (STR) in complex with tryptamine (PDB ID: 2FPB) was used as the initial model. All point mutations investigated in this study (V176F, M276L, and M276R) were introduced individually using PyMOL, followed by local backbone/side-chain optimization in Schrö dinger Protein Preparation Wizard. Hydrogen atoms were added according to physiological protonation states at pH 7.0, and crystallographic water molecules within the catalytic pocket were retained to preserve native hydrogen-bonding networks.

#### Reference Pose Generation for (S)-strictosidine

To obtain an experimentally consistent binding orientation, the structures of STR–tryptamine complexes (PDB IDs: 2FPB and 2FPC) were aligned. The bound conformation of (S)-strictosidine extracted from 2FPC was adopted as a reference pose. During docking, positional restraints (3 °A tolerance) were applied to maintain the substrate geometry relative to tryptamine.

#### Molecular Docking

Induced-fit docking (IFD) was performed in Schrödinger, allowing side-chain flexibility within 5 °A of the catalytic pocket. The docking grid box was generated using the size of the reference ligand (from 2FPC). Tryptamine was fixed in the binding site, while (S)-strictosidine was redocked using the constrained protocol.

#### Molecular Dynamics Simulations

Docking complexes were subjected to explicit solvent MD simulations using the AMBER99 force field for proteins and the GAFF force field for small molecules. The process included defining the box, solvating, energy minimization, heating, equilibration, and production. After neutralization by the addition of explicit counter ions (Na^+^ or Cl^−^) and energy minimization, a 500 ps NVT (constant temperature, constant volume) ensemble was performed to thermalize the systems to 300 K under constant volume and periodic boundary conditions. Then, a 500 ps NPT (constant temperature, constant pressure) ensemble was performed at an isotropic constant pressure of 1 bar and temperature of 300 K with a 2.0 ps time constant using the Parrinello-Rahman pressure coupling algorithm. Finally, a 100 ns MD simulation was carried out under periodic boundary conditions with an integration time step of 2 fs. Each mutation (V176F, M276L, M276R) was simulated in triplicate, starting from different random velocity seeds to ensure statistical robustness. RMSD, RMSF, and *R*_g_ analyses confirmed system equilibration before free-energy calculations.

#### Binding Free Energy Calculations

Binding free energy calculations were performed using `gmx MMPBSA` v1.6.2, which implements the Amber MM/PBSA protocol on GROMACS trajectories. The trajectories obtained from the 100 ns production MD simulations were analyzed using a single-trajectory approach, in which the protein, ligand, and complex coordinates were extracted from the same MD trajectory to minimize structural inconsistencies between states. Frames from 20–100 ns were used for MM/PBSA analysis. Given a 10 ps trajectory sampling rate, this corresponds to analyzing 800 evenly spaced snapshots, ensuring statistically robust free energy estimates. The ligand parameters were taken from the GAFF parameterization used during MD simulation (AM1-BCC charges). The bondi (mbondi2-like) PB radii set (option 4) was used for Poisson-Boltzmann calculations. All free energy calculations were performed at 298.15 K, consistent with the MD simulation temperature. MM/PBSA decomposes the binding free energy according to:

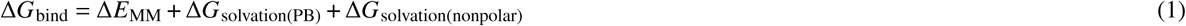

where Δ*E*_MM_ includes bonded, van der Waals, and electrostatic interactions; PB solvation energy is computed using the Poisson–Boltzmann equation; and the nonpolar contribution is estimated from the solvent-accessible surface area (SASA).

**Figure S4.**
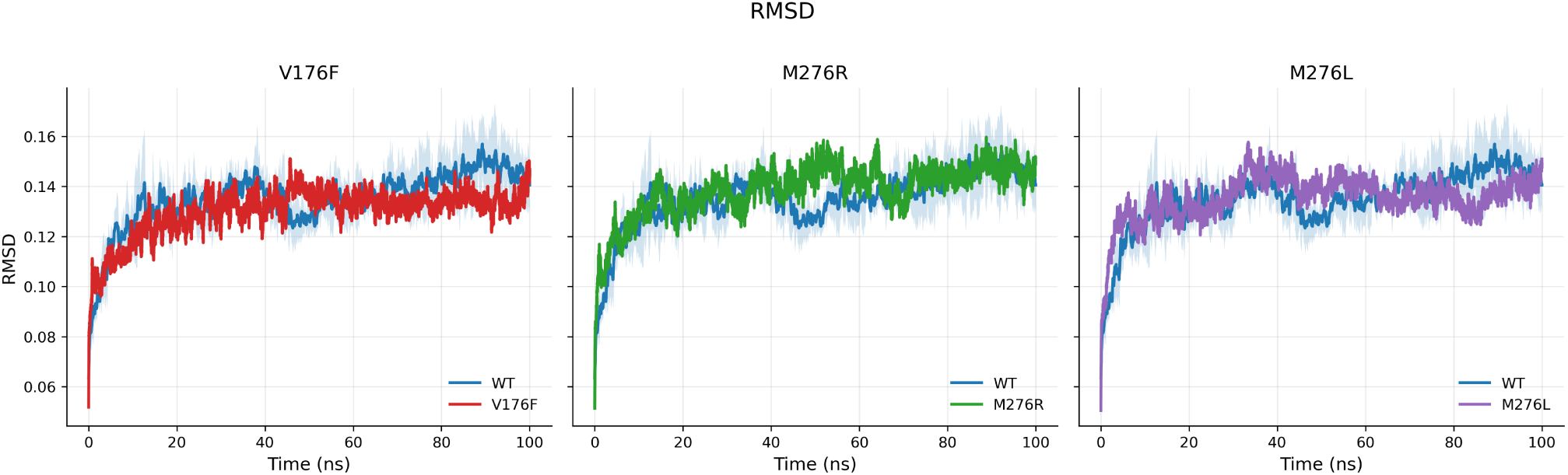
Backbone RMSD (ns) for WT versus variants (V176F, M276R, M276L). Each panel compares one variant to WT; solid lines show the mean across replicates and the shaded bands indicate the standard deviation. RMSD traces are lightly smoothed (21-point moving average) for visualization only.

**Figure S5.**
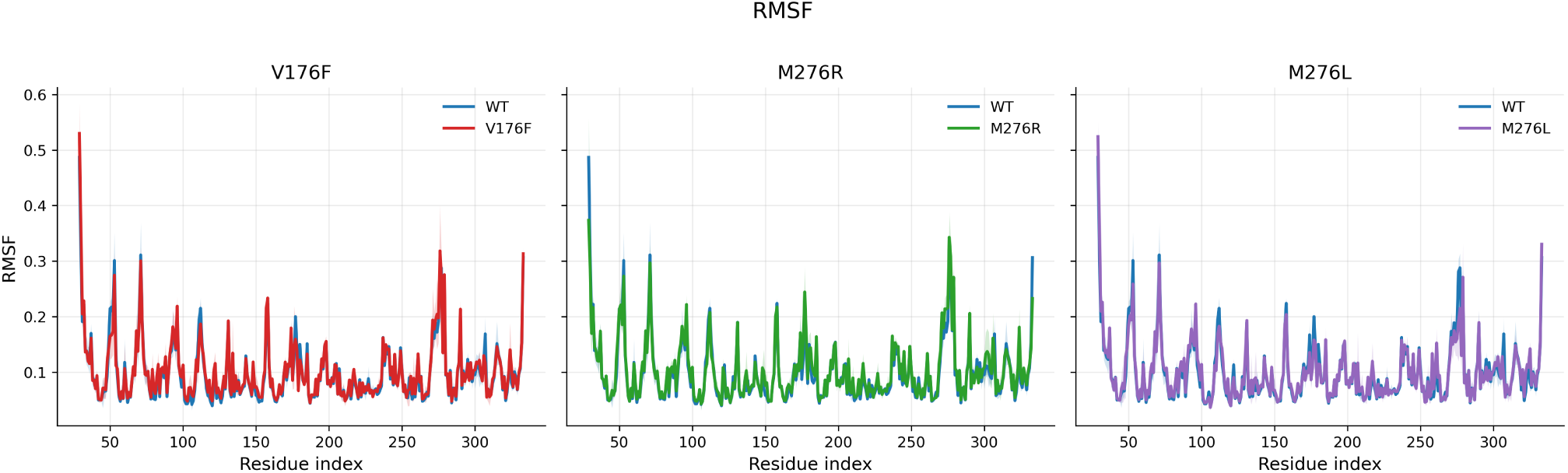
Per-residue RMSF for WT versus variants (V176F, M276R, M276L). Each panel compares one variant to WT; solid lines show the mean across replicates and the shaded bands indicate the standard deviation.

**Figure S6.**
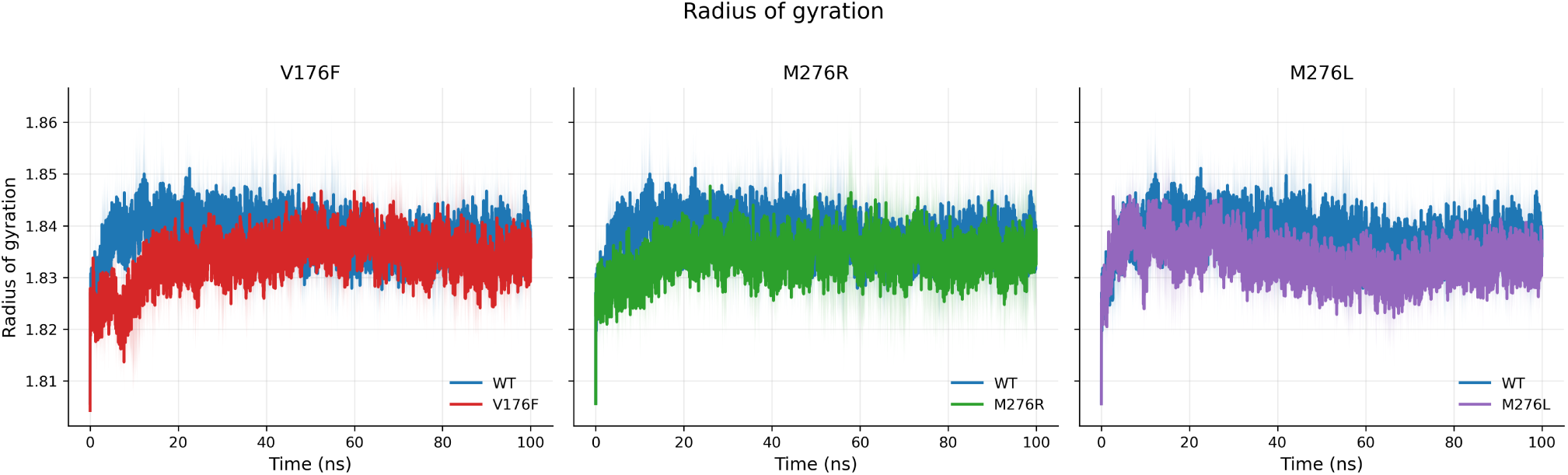
Radius of gyration (*R*_g_, ns axis on the trajectories) for WT versus variants (V176F, M276R, M276L). Each panel compares one variant to WT; solid lines show the mean across replicates and the shaded bands indicate the standard deviation.

#### Trajectory Analyses (RMSD, RMSF, and Radius of Gyration)

We assessed the stability and flexibility of wild-type (WT) and optimized variants by analyzing the protein backbone RMSD, per-residue RMSF, and the radius of gyration (*R*_g_) over production trajectories. For each system, replicate simulations were aggregated by computing the mean trace with a semi-transparent band indicating the standard deviation. Across systems, the backbone RMSD of the variants is comparable to that of WT, showing stable plateaus after equilibration and no evidence of excessive drift (Figure S4). Likewise, the per-residue RMSF profiles are broadly similar between variants and WT, with only minor, localized differences near engineered regions (Figure S5). By contrast, the radius of gyration is consistently lower for the variants than for WT (Figure S6), indicating a slightly more compact ensemble. Such modest compaction—together with preserved global stability—suggests reduced conformational heterogeneity and improved active-site pre-organization, providing a plausible dynamic basis for the higher catalytic activity observed for the variants relative to WT.

**Figure S7.**
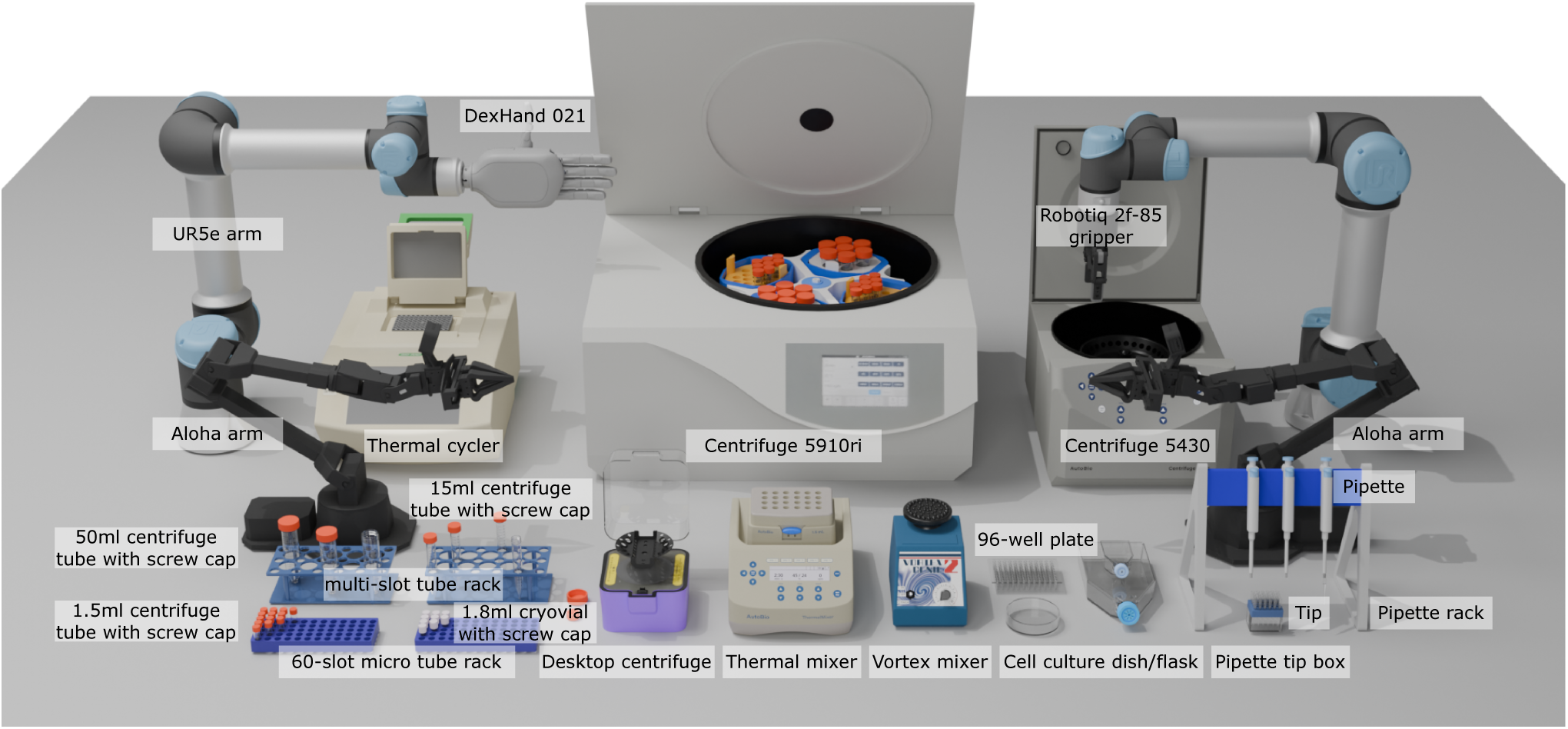
The overview of digital instruments and devices for virtual lab.

### VLA Training and Tasks

#### Simulation Environment and Data Preparation

To set up a high-fidelity environment for STELLA-VLA and baseline training, we extend asset modeling, physics simulation, and rendering capabilities specifically for biological primitives.

##### Asset Generation Pipeline

We construct dimensionally accurate digital models of laboratory assets (categorized into Instruments, Containers, Racks, and Robots) through a multi-stage workflow.

1. **Reconstruction:** We capture multi-view video of real-world instruments and utilize 3D Gaussian Splatting (3DGS) and the PGSR algorithm to reconstruct high-quality visual assets.
2. **Refinement:** Coarse meshes extracted from 3DGS are refined in CAD software to optimize topology for simulation while preserving critical geometric features and articulating joints.
3. **Standardization:** Refined models are UV-unwrapped with baked vertex colors and converted into the MJCF modeling language via a custom `gltf2mjcf` converter to define collision characteristics and physical properties.

##### Physics and Rendering Fidelity

Standard physics engines often lack the granularity required for lab equipment. We developed a suite of MuJoCo plugins to simulate specialized mechanisms prevalent in biology, including threads, detents, eccentric motion, and quasi-static liquid computation. Visually, we address the challenge of transparent materials (e.g., glassware) by integrating Blender’s Physically Based Rendering (PBR) pipeline, ensuring accurate refraction and reflection. Furthermore, we implement *dynamic texture rendering* for instrument screens, enabling the simulation of interactive digital UIs essential for VLA tasks.

#### Evaluation Tasks

We evaluate the model on four distinct biological manipulation tasks.

##### Scoring Metric

All tasks are scored binarily (1 for success, 0 for failure) with the exception of the *Operate thermal mixer panel* task. Due to the complexity of continuous value adjustments, this task utilizes a relative progress score to better reflect performance differences across policy iterations.

###### Transfer centrifuge tube

*Instruction:* “Pick up the centrifuge tube and move it to the other rack, row *{*target row*}*, column *{*target col*}*.” **Description:** Requires the robot (UR5e-Robotiq) to perform precise pick-and-place manipulation. The agent must visually identify the correct tube and interpret language instructions to locate a randomized target rack slot.

###### Operate thermal mixer panel

*Instruction:* “Adjust thermal mixer parameters, with speed set to *{*set rpm*}* rpm, temperature set to *{*set temp*}* ^◦^C, and time set to *{*set time*}* seconds.”

**Description:** Involves fine-grained interaction with a digital interface. The agent must read the current UI state via vision, interpret the target values from the instruction, and manipulate the touchscreen to adjust time, temperature, and frequency. Parameters are randomized to test cross-modal numerical reasoning.

###### Load centrifuge rotor

*Instruction:* “Insert a second centrifuge tube into the slot that is symmetrically opposite to the currently placed tube.” **Description:** Tests advanced geometric reasoning and precise positioning. The agent must perceive the location of a pre-existing tube within a centrifuge rotor and load a new tube into the symmetrically opposite slot to ensure balance. Both the rotor angle and the initial tube position are randomized.

###### Aspirate with pipette

*Instruction:* “Dual-UR5e pipetting: one arm lifts centrifuge tube, the other aligns pipette tip and aspirates liquid.” **Description:** A bi-manual task requiring high-level coordination between a holder arm (UR5e-Robotiq) and an actuator arm (UR5e-DexHand). The agent must stabilize the tube and precisely align the pipette tip to aspirate a specific volume of liquid. Liquid volumes are randomized to evaluate visual liquid-level sensing capabilities.

##### STELLA Training Algorithm

The training pipeline for STELLA is formalized in Algorithm 1. We denote the VLA policy as *π*_*θ*_, parameterized by *θ*, and initialize an empty replay buffer D. For a high-level task instruction *T*, the system first generates a sequence of interpretable subtasks *S* = {*s*_1_, …, *s*_*N*_ }. During the execution of each subtask *s*_*i*_, the policy generates a rollout trajectory *τ*_*rollout*_. A set of multimodal verifier tools, denoted as V, acts as the monitoring mechanism to detect execution failures. Upon detection of an error, the system interrupts the nominal policy and queries the recovery tools R to select an appropriate recovery policy *π*_*rec*_. If the resulting corrective trajectory *τ*_*rec*_ successfully resolves the error, it is aggregated into D alongside successful nominal rollouts. The policy parameters *θ* are then updated via supervised fine-tuning on the enriched dataset D, allowing the VLA to iteratively internalize robust recovery behaviors.

###### Algorithm 1 STELLA: Iterative Decompose-Monitor-Recover VLA Training

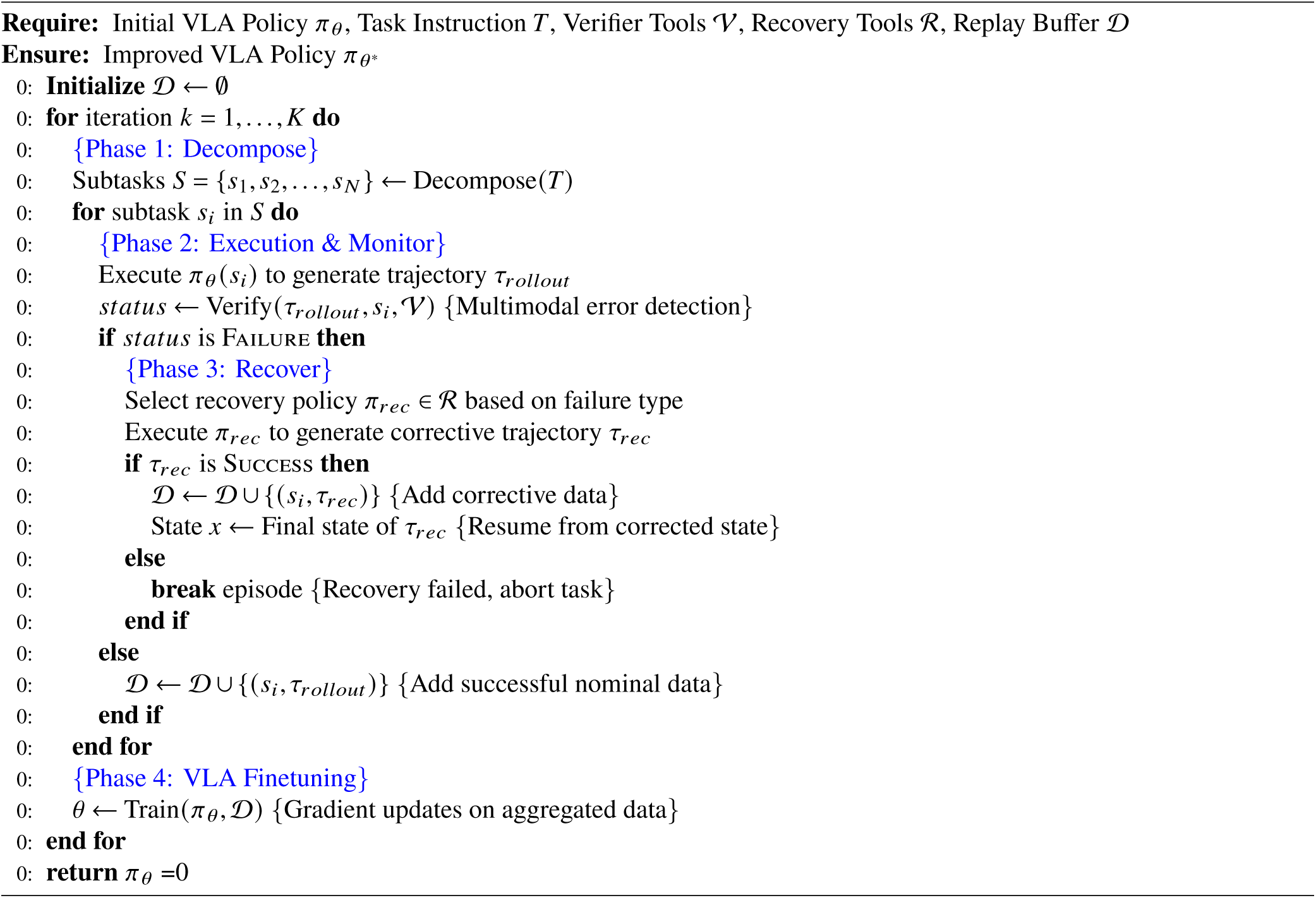

## Notes

### Competing Interest Statement

The authors have declared no competing interest.

### Summary of Updates

Added experimental verifications, such as figure 3, 4, 5

